# Identification of Potential Regulatory Non-Coding RNAs in *Lotus Japonicus* Symbiosis

**DOI:** 10.64898/2026.05.19.726297

**Authors:** Asa Budnick, Delecia Utley, Zuzanna Blahovska, Simona Radutoiu, Heike Sederoff

**Affiliations:** Department of Plant and Microbial Biology, North Carolina State University, Raleigh, NC, USA; Department of Molecular Biology and Genetics, Aarhus University, Aarhus, Denmark

**Keywords:** *Lotus japonicus*, Rhizobia, symbiosis, non-coding RNA, circular RNA, microRNA, gene expression, backsplice-generated miRNA response element (bsg-MRE)

## Abstract

Symbiosis between legumes and rhizobia is beneficial on nutrient-poor soils, as it enables the fixation of atmospheric N_2_. To establish this symbiosis, gene expression in both the host plant and the symbiont has to be regulated. To understand the underlying RNA-mediated regulation of host gene expression, we designed experiments to identify competing endogenous networks involving circular RNA, microRNA, and linear transcripts during symbiosis, using wt and symbiosis-deficient *Lotus japonicus* mutants with the rhizobium *Mesorhizobium loti (M. loti*).
CircRNA, miRNA, and linear transcripts were identified from *Lotus japonicus* wildtype and CCamK mutant (*ccamk-13; snf-1*) seedlings without inoculation or with *M. loti* inoculation using deep short-read sequencing with rRNA-depletion and random primers.
Differentially expressed miRNAs showed negative correlations to predicted target genes and may regulate symbiotic processes. The symbiosis essential iron-sensor *LjnsRING/BRUTUS* expresses a circRNA which was upregulated in symbiotic treatments. This circRNA may act as a target mimic and contribute to nodule longevity. CircRNAs are predicted to act predominantly as *trans-*regulatory molecules with similar frequencies in *Arabidopsis thaliania, Oryza sativa,* and *Lotus japonicus*.
We identified novel miRNAs, long noncoding RNAs, and circRNAs, and nominated several as potential new regulatory non-coding RNAs that may act as target mimics to stabilize genes and support symbiosis.

**Summary:** Symbiosis between *Lotus japonicus* and *Mesorhizobium loti* involves treatment-specific regulation of competing endogenous RNA networks involving circular RNA, miRNA, and linear transcripts.

## INTRODUCTION

Symbiosis between legume hosts and rhizobial bacteria can result in mutual beneficial N_2_ assimilation under low soil nitrogen conditions. To establish symbiosis, both plants and bacteria undergo multiple developmental stages, from signaling and nodule formation to functional N_2_ fixation and metabolite exchange. These programs require coordinated changes in gene expression for both the plant host and bacterial symbiont. These changes in gene expression are not only locally regulated at the site of infection and nodulation but also require systemic regulation to adjust processes of photosynthetic carbon assimilation and nitrogen use, thereby affecting overall plant growth and development (Lepetit & Brouquisse 2023).

Different forms of non-coding RNA, including microRNAs (miRNAs), long-noncoding RNAs (lncRNAs), and circular RNAs (circRNAs), have been shown to regulate gene expression at transcriptional and translational levels, locally and systemically (Wang *et al*. 2019). MiRNAs are short (21-24nt) RNAs that are expressed as nuclear genes and processed as RNA effectors of post-transcriptional silencing via degradation and translational inhibition. During nodulation, miRNA2111 is a key mobile component in the Autoregulation Of Nodulation (AON) pathway to control nodule number based on plant need (Tsikou *et al*. 2018).

Long non-coding RNAs (lncRNA) are transcripts larger than 200nt without a large open reading frame. In rhizobial symbiosis the lncRNA ENOD40 is highly expressed in nodules and suppression of ENOD40 reduces nodule number in *Lotus japonicus* and *Medicago truncatula* (Yang *et al*. 1993; Campalans *et al*. 2004; Kumagai *et al*. 2006). LncRNAs can serve as target mimics - harboring miRNA response elements (MREs) which can sequester miRNAs and stabilize the expression of other miRNA targets (Wu *et al*. 2013). Recently, ENOD40 was described as a target mimic for miR169 in *Medicago truncatula* which stabilizes expression of NF-YA1 and supports nodulation (Wijsman *et al*. 2025).

Circular RNAs are single-stranded covalently closed non-coding RNAs produced via back-splicing of transcripts where the 3’ and 5’ ends of a portion of a transcript are ligated to form the back-splice junction (BSJ) (Zhang *et al*. 2024). CircRNAs are emerging as gene regulatory molecules in plants. CircRNAs can form R-loops and influence alternative splicing of transcripts from their locus of origin. In *Arabidopsis thaliana* overexpression of a circRNA from exon 6 of the *SEPALLATA3* gene resulted in increased R-loop formation, a larger proportion of the exon-6-skipped alternative splice isoform, and an increase in petal count (Conn *et al*. 2017). CircRNAs can also act as competing endogenous RNAs (ceRNAs) to bind miRNAs and stabilize the expression of other targets. Knockout of a circRNA in rice bearing MREs for OsMIR408 led to increased abundance of OsMIR408 and decreased abundance of 7 of 9 measured OsMIR408 targets (Zhou *et al*. 2021). Studies have identified differentially expressed circRNAs during symbiotic nitrogen fixation in *Lotus japonicus, Glycine max, Medicago truncatula,* and *Phaseolus vulgaris* (Wu *et al*. 2020; Jing *et al*. 2022; Lin *et al*. 2025; Utley *et al*. 2026).

In this study we investigate the hypothesis that non-coding RNAs contribute to the regulation of rhizobial symbiosis via ceRNA networks. We identified mRNAs, miRNAs, lncRNAs, and circRNAs expressed in *Lotus japonicus* plants after treatments with and without rhizobial symbiont and identified potential ceRNA networks. We identified non-coding RNAs that are upregulated in symbiosis treatments and are connected by predicted MREs to genes with known or potential roles in symbiosis. In particular we identified a circRNA expressed from the rhizobial symbiosis-essential *LjnsRING/*BRUTUS gene that may act as a target mimic and indicate that BRUTUS has both coding and non-coding transcripts that could play a role in the regulation of symbiotic nitrogen fixation (Shimomura *et al*. 2006). We also introduce a novel classification of circRNAs as *cis-* or *trans*-regulatory molecules and find evidence of conservation of circRNA parent genes and *trans-*regulatory circRNAs across 6 and 3 plant species respectively.

## MATERIALS AND METHODS

### Plant growth, bacterial culture, and inoculation

*Lotus japonicus* seeds (Gifu B-129, *snf-1, ccamk-13*) were surface-sterilized in 1% (v/v) sodium hypochlorite (15 min), washed five times in sterile water, and imbibed overnight at 4°C in the dark. Seeds were stratified on wet sterile filter paper for germination under 16h light (21°C) and 8h dark (19°C). After seven days, seedlings were transferred to plates containing Agar Noble with 0.25X Broughton & Dilworth (B&D) medium, and the plates were covered with filter paper (Broughton & Dilworth 1971). *Mesorhizobium loti* (R7A) was cultured for 2 days at 28°C in yeast mannitol broth (YMB). The 9-day-old seedlings were inoculated with 750 µL of a bacterial suspension (OD₆₀₀ = 0.02) along the length of the roots. Plants were grown for another 3- or 5-weeks post-inoculation. Root tissue was harvested into liquid N_2_ and stored at - 80℃.

Prior to harvesting, all plates were imaged. Root lengths and shoot heights were measured in FIJI (ImageJ). Pink and white nodules were counted per plant.

### RNA extraction and sequencing

Total RNA was extracted from 50–100 mg of ground *L. japonicus* root tissue using a modified CTAB protocol (Jordan-Thaden et al., 2015) followed by TRIzol purification (for details see Suppl. File 1). Each sample represented RNA pooled from 15 plants.

Sequencing libraries were prepared separately for the sequencing of small RNA (50bp-PE) and ribosomal RNA-depleted total RNA (150bp-PE) using the Illumina platform. Reads were quality checked with FastQC (Andrews, 2010). After QC, the bioinformatic pipelines are distinct for the 150bp-PE and 50bp-PE sequencing data.

### Analysis of linear coding and non-coding RNA

An overview graphic and detailed methods for bioinformatics pipelines used here are provided in Suppl. File 1. In short, 150bp-PE reads were trimmed with Trimmomatic (v0.39) in paired-end mode. Linear RNA was aligned to the two different genomes *Lotus japonicus* Gifu v1.3 and *Mesorhizobium loti* R7A (GCA_012913625.1_ASM1291362v1) using BBsplit (v38.90) with a minimum identity set to 97% FeatureCounts (v2.0.1) was used to quantify the number of fragments mapping to annotated genes for *M. loti* (Bushnell 2014; Liao *et al*. 2014; Bolger *et al*. 2014). Custom R scripts were used for further data analysis.

For analysis of *L. japonicus* linear RNA expression, trimmed reads were mapped to the *L. japonicus* genome with Hisat2 (v2.1.0) with parameters --pen-noncansplice 4 and --rna-strandness RF (Kim *et al*. 2019). Hisat2 alignments were used to create and merge isoform-level annotations using stringtie (v3.0.0) with strandedness flag -rf and flag -N for nascent transcript processing (Shinder *et al*. 2025). Transcript sequences were pulled from the stringtie assembly with gffread (v1.0) (Pertea & Pertea 2020). IsoformSwitchAnalyzeR was used in conjunction with custom R scripts for isoform and differentially expressed gene analysis alongside EdgeR, and DEseq2 (Robinson *et al*. 2010; Love *et al*. 2014; Vitting-Seerup & Sandelin 2019). Transcripts were categorized as long non-coding RNAs if they were between 200 and 10,000 nt, had a coding potential of less than 0.5 by CPC2 (V2), and did not have a strong overlap with a single coding gene (Kang *et al*. 2017).

### CircRNA analysis

CircRNAs were identified using two pipelines, CLEAR (v1.0.1) and CIRI-full (v2.0) (Zheng, Ji, *et al*. 2019; Ma *et al*. 2019). CIRI-full requires untrimmed reads and generates circRNA identities as well as full sequences of circRNAs where there is sufficient sequence information. CIRI-full was run with default parameters recommended in the CIRI-Cookbook (Zhang 2021). Trimmed sequences were input into CLEAR, which identifies circRNAs and produces estimates of linear and circular RNA expression. CircRNA information from CLEAR and CIRI-full was merged with custom R scripts.

### PCR validation of select circRNAs

Validations of select circRNAs were performed with RNA treated with RNase R to digest linear RNAs. RNA (∼2 µg) was treated with 20 U RNase R (Lucigen) in a modified LiCl buffer according to Xiao and Wilusz (2019) or treated in the same buffer and incubation conditions without the addition of RNase R enzyme (RNase R -) (Xiao & Wilusz 2019). RNA reactions were purified with 1.8× SPRI, quantified using the Qubit High Sensitivity RNA kit (Invitrogen), and used for cDNA synthesis. cDNA synthesis was performed using DNase-treated total RNA and SuperScript IV reverse transcriptase with random hexamer primers (Thermo Fisher Scientific), following the manufacturer’s instructions. Divergent primers were designed to span the back-splice junction (BSJ) and maximize internal circRNA sequence coverage. PCR amplification was performed using OneTaq Hot Start 2X Master Mix (New England Biolabs) on cDNA (10% of total volume) and on 100 ng of RNase A-treated genomic DNA (gDNA) as a control for false positives. PCR conditions and primers are available in supplementary file 2.

Amplicons were visualized via agarose gel electrophoresis. PCR products of expected size were gel-excised, purified, and submitted for Sanger sequencing. Strand-specific cDNA synthesis was performed with single primers to validate the strandedness of RNAs from the BRUTUS locus, following cDNA synthesis PCR conditions were the same as above.

### CircRNA Conservation Analysis

Previously reported circRNAs were obtained from PlantCircRNA for *Medicago truncatula* and *Glycine max*. CircRNAs and full sequences were obtained from Chu *et al*. 2022 for *Arabidopsis thaliana* and *Oryza sativa* (Chu *et al*. 2022). CircRNAs from *Phaseolus vulgaris* were obtained from Wu *et al*. 2020 (Wu *et al*. 2020). Protein sequences were obtained for 8 plant species: *Amborella Trichopoda* (Amborella_trichopoda.AMTR1.0.pep.all) (EnsemblPlants, v62), *Arabidopsis thaliana* (Arabidopsis_thaliana.TAIR10.pep.all) (EnsemblPlants, v62), *Glycine max* (Glycine_max.Glycine_max_v2.1.pep.all) (EnsemblPlants, v62), *Lotus japonicus* (Lotus_japonicus_Gifu_v1.3) (Lotus Base), *Medicago truncatula* (Medicago_truncatula.MtrunA17r5.0_ANR.pep.all) (EnsemblPlants, v62), *Oryza sativa* (Oryza_sativa.IRGSP-1.0.pep.all) (EnsemblPlants, v62), *Phaseolus vulgaris* (Pvulgaris_442_v2.1.protein) (DOE-JGI, Phytozome 14), *Physcomitrium patens* (Physcomitrium_patens.Phypa_V3.pep.all) (EnsemblPlants, v62) (Mun *et al*. 2016; Yates *et al*. 2022). Identification of orthogroups was performed using OrthoFinder (v3.1.2) with default parameters (Emms *et al*. 2025). Orthogroup and circRNA-parent orthogroups were analyzed with custom R scripts. The number of orthologous parent genes who produced circRNAs across species intersections was tallied and compared to a set of 100 samplings of randomly assigned circRNA-parents, which contained the same proportion of circRNA parent orthogroups across each species.

### MicroRNA analysis

Version 1.1.2 of sRNAminer was used with default parameters to identify loci for miRNAs from read 1 and read 2 of 50bp-PE reads (Li et al., 2024). Cleaning of ribosomal RNA, organellar RNAs, and other non-coding RNAs not of interest for this analysis was done using annotated non-coding and organellar sequences from *Lotus japonicus* (Genome version LjGifuV1.2 (Kamal et al., 2020)). Output annotation files were merged using Another Gene Annotation Toolkit (v1.4) (Dainat et al., 2020).

To take advantage of PE sequencing and get more accurate counts, read 1 and read 2 were adapter-trimmed and merged using bbduk and bbmerge (BBMap v38.86) and then collapsed into counts using sRNAminer seq-collapse. Collapsed counts were aligned to the Lotus genome using bowtie, and then miRNA expression was estimated using sRNAminer bowtie-to-sRNAminer and miRNA_abundance functions. SRNAminer outputs a combination of -3p, -5p, and -mature, -star depending on its matching loci to and abundance (the more abundant mature miRNA is given the -mature label and the other the -star). Custom R scripts were used for further analysis and to determine the sequences of the final set of mature miRNAs.

### miRNA Target Prediction and Conserved Targeting Analysis

Raw fastq files were processed with CIRI-full (v2.0) pipeline (Zheng & Zhao, 2018) to assemble full-length circRNA sequences. To test for back-splice generated miRNA response elements (bsgMREs), circRNA sequences were represented both as the original full sequence from CIRI-full and as an offset version of that sequence made by repositioning the last 40 nt to the sequence start. Both original and ‘offset’ circRNA sequences were analyzed for predicted miRNA response elements (MREs) using psRNATargetV2 (Dai & Zhao, 2011) with miRNAs from sRNAminer. In cases where the ‘offset’ circRNA sequences were predicted to have distinct MREs compared to the original sequence, these new MREs were termed bsgMREs.

CircRNA sequences from *Arabidopsis thaliana* and *Oryza sativa* were queried in the same manner using psRNATargetV2 against miRNA sequences from miRbase (Kozomara & Griffiths-Jones 2014). Mature miRNA sequences from *Lj, At,* and *Os* were clustered in R using DECIPHER (v3.2.0) with between 3-10 iterations and a cutoff of 0.3 (Wright 2024). Conserved targeting of circRNAs was determined by a shared miRNA cluster having predicted circRNA targets with parent genes in the same orthogroup.

All custom R scripts are available at https://github.com/SederoffLab/Lotus_Cousins_Friends.

## RESULTS

### Physiological analysis

We grew *L. japonicus* wild type Gifu plants as well as two mutant versions of the symbiosis essential Calcium/Calmodulin-dependent protein kinase gene (CCaMK) (*LotjaGi3g1v0307700*) on media without nitrogen. Plants were either mock or inoculated with *Mesorhizobium loti* (R7A) and harvested after 3 or 5 weeks (wpi). The *snf-1* mutant version of CCaMK contains a Threonine to Isoleucine substitution (T265I) in the CCaMK kinase domain, resulting in an autoactive CCaMK isoform that promotes spontaneous nodulation in the absence of rhizobia. The *ccamk-13* mutant cannot form nodules even in the presence of rhizobia due to a 7bp insertion, which results in a premature stop codon and no active CCaMK expression (Tirichine *et al*. 2006; Singh & Parniske 2013). Mutant sequences were confirmed in RNAseq data (Fig.S1). An overview of treatments is shown in Fig. 1a. Treatment conditions are abbreviated as genotype (G=gifu wt, C=*ccamk-13*, S=*snf-1*), number of weeks post inoculation, and inoculation condition (R=*Mesorhizobium loti* inoculation, N=mock inoculation); for example, G3R indicates wt plants were grown for three weeks post inoculation with *Mesorhizobium*.

**Fig. 1.**
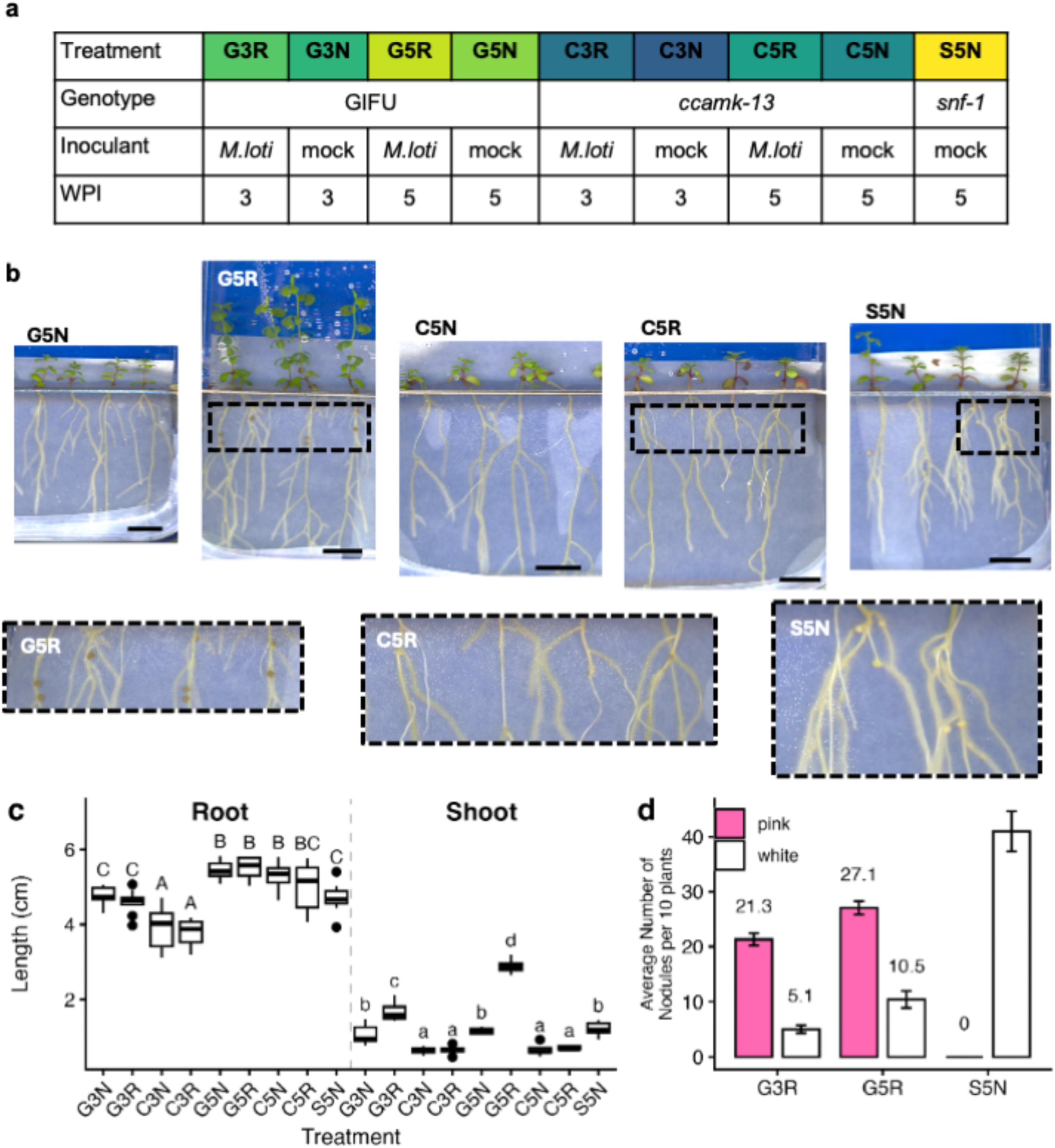
Plant growth and nodule development. a) Treatment overview. Three genotypes were inoculated with either M. loti or a mock solution and incubated for 3 or 5 weeks before harvesting for physiological measurements and RNA sequencing. The first letter in a treatment acronym identifies the genotype (G, Gifu; C, *ccamk-13*; S, *snf-1*), while the second letter refers to weeks after nodulation (3wpi, 5wpi) and the last letter indicates whether the roots were inoculated with rhizobia (R) or mock inoculant (N). b) Representative 5wpi plants and close-up of select roots. Representative images from 3-week treatments are in Fig. S2. c) Primary root length and shoot height, letters indicate statistically significant groups (Pairwise Wilcoxon Rank Test, n=90, FDR <0.05). d) Average number of pink and white nodules per 10 plants (N=9 observations of 10 plants).

Lotus seedlings grown on N-free medium showed the expected growth differences in response to time, genotype, and inoculation (Fig. 1b, Fig. S2). We measured primary root length, shoot height, and the number of nodules that either appear pink/brown due to leghemoglobin, and white nodules, which do not contain active bacteroids (Fig. 1c, d). There was no difference in root length between inoculated and non-inoculated seedlings within a genotype (Fig. 1c). The roots of *ccamk-13* seedlings were shorter than those of Gifu seedlings at 3 wpi independent of inoculation. Non-inoculated *snf-1* roots were shorter than Gifu or *ccamk-13* roots at 5 wpi. This suggests that nodule development without symbiotic nitrogen fixation, as observed in the *snf-1* mutant, results in shorter roots.

Shoot height was most affected by genotype and inoculation (Fig. 1c). At 3 and 5 wpi the shoots of Gifu seedlings inoculated with *M. loti* were taller than uninoculated treatments. These two treatments (G3R, G5R) are the only ones with pink/brown nodules capable of fixing N_2_ and providing the plants with nitrogen to support growth. The *ccamk-13* plants had shorter shoots than all other treatments whereas *snf-1* plants had similar shoot growth to non-inoculated wt plants. Qualitatively, the 5wpi non-symbiotic treatment plants showed visible stress phenotypes, such as purple leaves and stems (Fig 1b, Fig. S2.).

G5R plants showed a higher percentage of mean inactive nodules out of the mean total nodules (10.5/37.6 or 28% per 10 plants) compared to G3R (5.1/26.4 or 19% per 10 plants) (Fig. 1d), likely indicating a second round of nodulation had occurred in G5R plants but these younger nodules were not yet active.

### Deep sequencing with random primers yields lotus and rhizobial transcriptomes

Identification of circRNA via short-read sequencing requires the use of random primers because circRNAs do not have poly-A tails that are usually targeted in transcriptome sequencing. The transcriptome generated using random primers includes circRNAs and diverse classes of linear RNA, including mRNAs and non-coding RNAs. Sequencing yielded an average of 204 million reads per sample mapped to *Lotus japonicus* (Fig. S3).

Significantly fewer reads mapped to the *M. loti* genome in the inoculated *ccamk-13* samples compared to the inoculated Wt samples (Fig. S3). Most differentially expressed *M. loti* genes were up-regulated in Wt versus *ccamk-13* plants (Fig. S4a, supplemental file 3). Nitrogenase and respiration genes associated with symbiotic nitrogen fixation were more highly expressed in symbiosis treatments. Transcripts involved with mobility, like flagellar proteins and nitrate uptake and reduction, were more highly expressed in rhizobia inoculated on *ccamk-13* mutants compared to those inoculated on Gifu plants (Fig S4b). This is consistent with ‘free-living’ rhizobia metabolism in *ccamk-13* mutants.

### Distinct RNA classes respond differently to symbiosis treatment conditions

Expression profiles of linear mRNAs, miRNAs, and circRNAs showed distinct and class-specific responses to symbiosis-related treatment conditions (Fig. 2). Linear mRNA showed separation along symbiosis and weeks-post-inoculation axes (Fig. 2a). MiRNA expression showed a similar structure, with symbiotically active samples being distinct from all other treatments; however, separation between G3R and G5R was less pronounced relative to mRNAs (Fig. 2b).

**Fig. 2.**
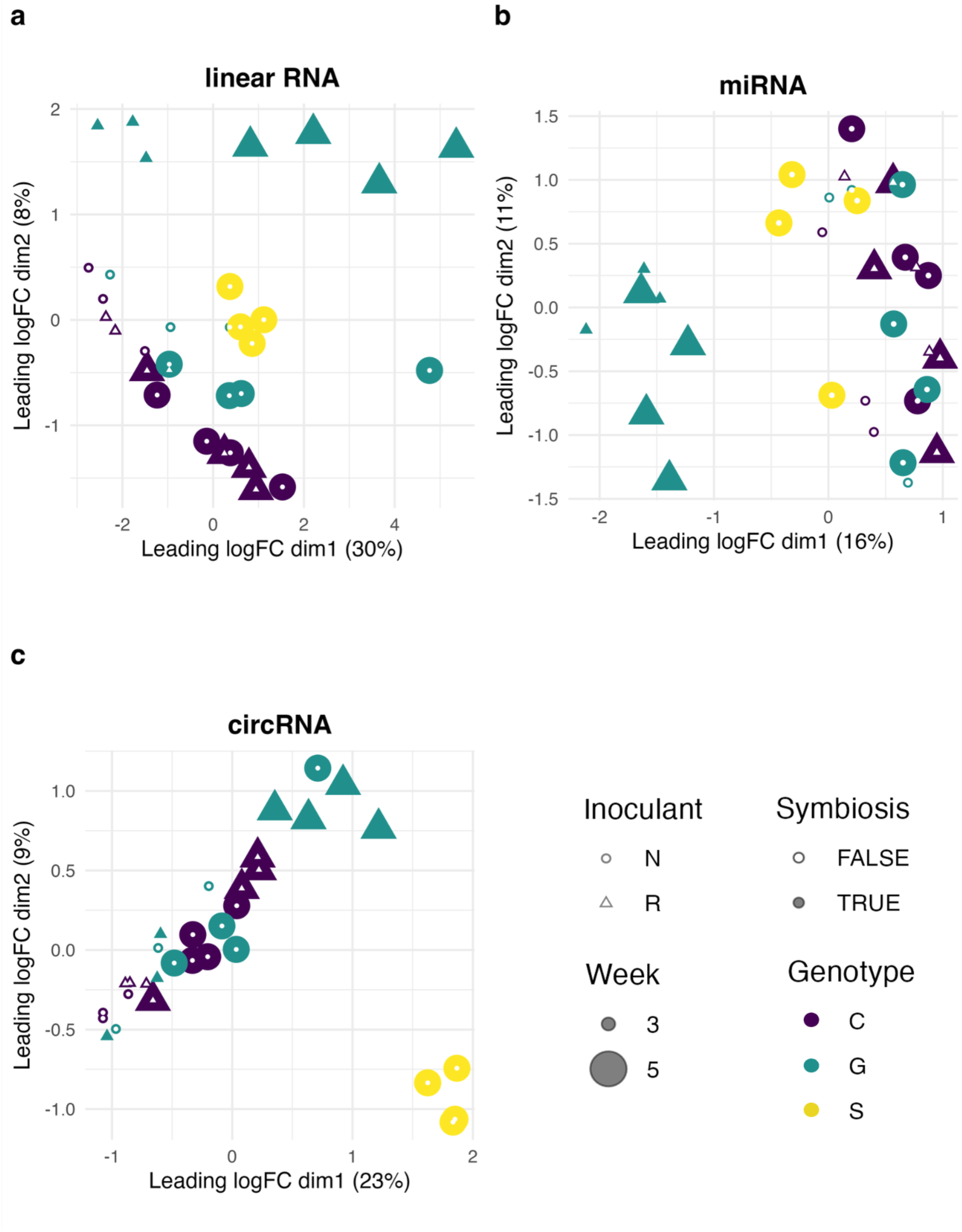
Multi-dimensional scaling of 3 classes of RNA. a) Linear RNA transcript expression shows separation of dimension 1 according to weeks post inoculation and dimension 2 based on active symbiosis. b) Micro RNA expression shows clustering along dimension 1 by active symbiosis. c) *snf-1* mutant plants show a distinct cluster of Circular RNA expression whereas treatment wpi and active symbiosis do not show distinct clusters.

In contrast to linear mRNAs and miRNAs, circRNA expression was structured by genotype (Fig. 2c). The *snf-1* mutant showed pronounced separation from other genotypes, indicating a strong association between plant genotype and circRNA abundance. This separation was driven by differences across hundreds of circRNAs rather than a small set of specific circRNAs and we were unable to identify an explanatory pattern between snf-1 responsive circRNAs and coding transcripts. *Ccamk-13* and Gifu <u>circRNA expression also</u> clustered along a weeks-post-inoculation axis. These patterns highlight the distinct expression profiles among three different RNA classes under the same experimental conditions.

### Coding and non-coding linear RNAs respond to active symbiosis

Pairwise comparison of treatments showed 27,193 genes were differentially expressed (FDR < 0.01, Absolute Log_2_FC > 1) in at least one treatment comparison. Clustering of the top 5000 DEGs by minimum FDR across all pairwise treatment comparisons shows distinct clusters of DEGs responding to symbiosis and genotype (Fig. 3a, supplementary file 4). Symbiosis up-regulated clusters include known key transcriptional regulators of nodule organogenesis like NODULE INCEPTION (NIN), NIN-LIKE PROTEINS (NLPs) and GLUTAMYL-TRNA REDUCTASE involved in regulating heme and leghemoglobin biosynthesis, as well as TOO MUCH LOVE (TML), a nodulation repressor that is the target of a systemic miRNA2111 regulating shoot-controlled root nodulation (Tsikou *et al*. 2018; Wang *et al*. 2025). Homologs of the key regulators of iron status in Arabidopsis, the zinc-finger containing E3 ligase BRUTUS (LotjaGi4g1v0453800) and the bHLH transcription factor POPEYE (LotjaGi4g1v0248800) are also in the symbiosis up-regulated gene clusters (Fig. 3b) (Long *et al*. 2010; Hindt *et al*. 2017). Gene ontology analysis showed that terms related to glycolysis, nitrate response, and amino acid transport were enriched in symbiosis upregulated clusters (Fig. 3c)

**Fig. 3.**
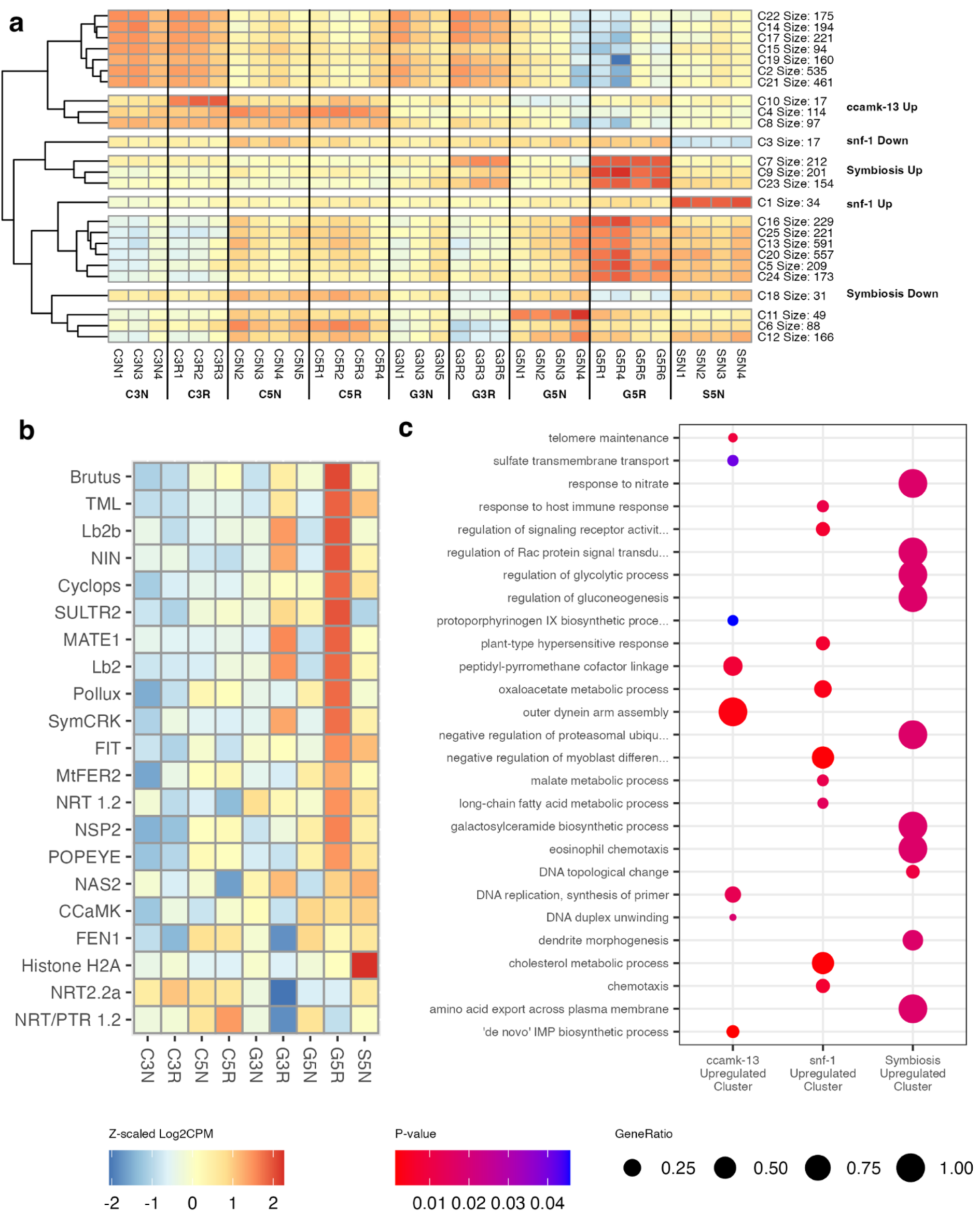
Differential gene expression. a) K-means clustering of Z-scaled log2cpm for the top 5000 DEGs by lowest FDR. Complete data set in supplementary file 4. b) Relative expression (Z-scaled log2cpm) of select genes involved in the Common Symbiosis Pathway, nodule organogenesis, iron and hemoglobin metabolism or otherwise of interest. c) Biological Process GO term overrepresentation analysis of select labeled clusters of plant gene expression.

Similar clustering of transcript-level expression revealed clusters upregulated in symbiotically active treatments as well as a cluster upregulated in *ccamk-13* mutant treatments (Fig. S6). The symbiosis upregulated clusters include 16 potential non-coding RNAs (Fig. S7). Among these is LotjaGi2v1ncRNA9, which overlaps with the known symbiosis-associated lncRNA ENOD40 (MSTRG.10902.1; LjG1.1_chr2:79764218-79764926). Two symbiosis upregulated potential novel non-coding RNAs can be classified as natural antisense RNAs. LotjaGi1v1ncRNA14 (MSTRG.5920.5) is antisense to LotjaGi1g1v0576500, a putative universal stress protein. The expression of LotjaGi1v1ncRNA14 and LotjaGi1g1v0576500 shows a weak positive correlation (Spearman Rank Test, p-value = 0.016, R^2^=0.18, Fig. S8a). LotjaGi1v1ncRNA23 is upstream and antisense of LotjaGi1g1v0785900 a putative F-box containing protein, these are also positively correlated (Spearman Rank Test, p-value < 1x10^-6^, R^2^=0.62, Fig. S8b).

### Novel differentially expressed miRNAs may regulate symbiosis-associated genes

Small RNA sequencing identified 502 unique mature miRNAs, of which 301 had at least 10 total reads across all samples and were included in differential expression analysis (supplemental file 5). MiRNAs without matches in reference databases (miRBase, PmiREN, or sRNAanno) were considered novel, resulting in the identification of 64 novel miRNAs (Kozomara & Griffiths-Jones 2014; Chen *et al*. 2021; Guo *et al*. 2022). Among the known miRNAs, several families were highly represented with multiple distinct miRNA genes, including miR399, miR166, miR319, miR395, miR172, and miR2111 (Fig. 4a, Fig. S8a).

**Fig. 4.**
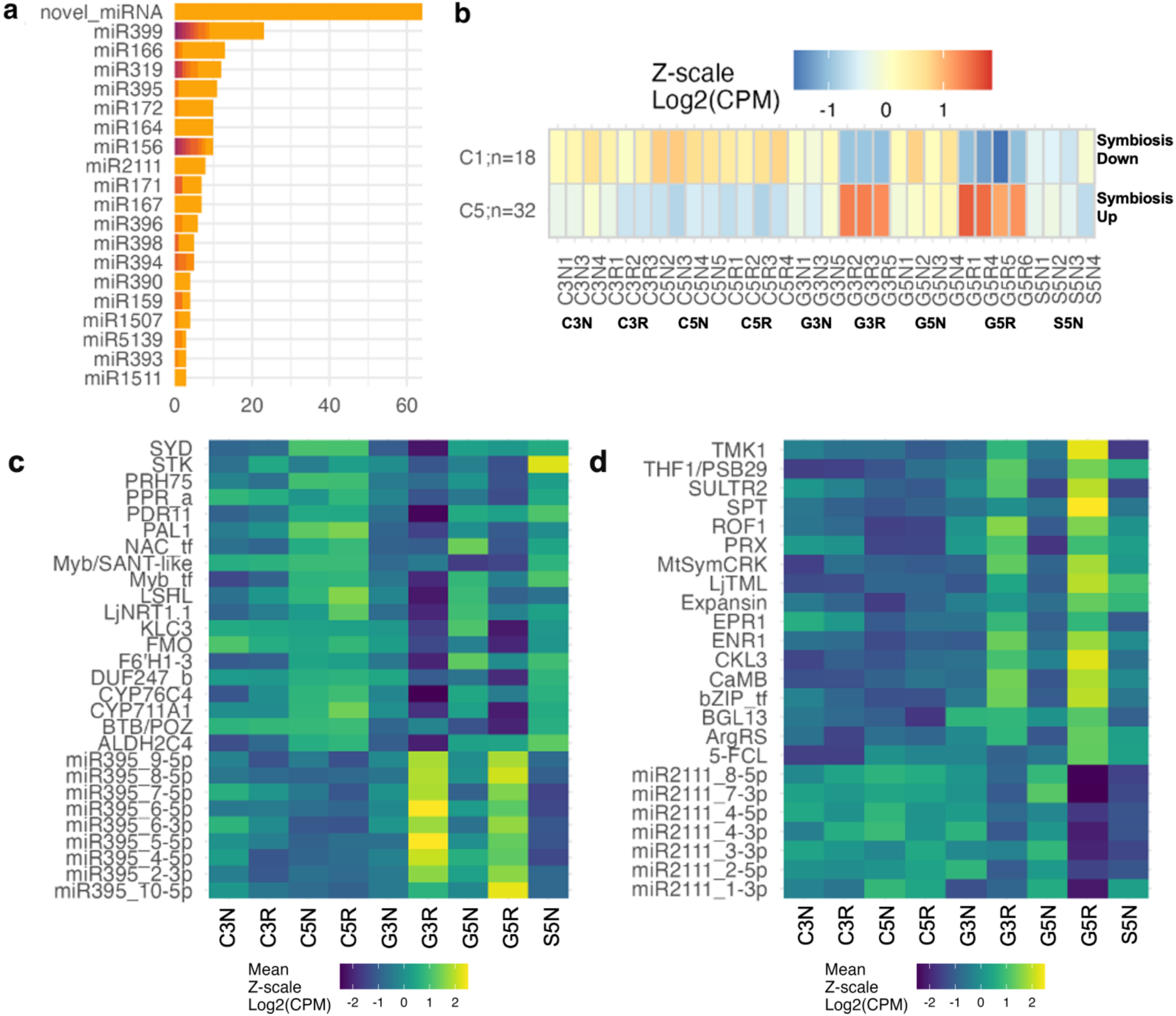
Diversity and expression patterns of miRNAs. a) Number of miRNA genes identified across different miRNA families, a color gradient is assigned to each lettered version of a miRNA gene identified (i.e. miR156c, miR156d, etc.) as well as the unlettered version (i.e. miR156). Only miRNA families with more than 2 members are shown here; the full version of the plot is available in Fig. S8a. b) Heatmap of k-means clusters of miRNA expression highlighting 2 clusters that respond to active symbiosis. Full heatmap in Fig.S8b. Expression of miR395 (c) and miR2111 miRNAs which show symbiosis dependent regulation and negative correlations with predicted target genes (Pearsons, FDR <0.05). Expression is shown as treatment average Z-scaled log2(CPM) of miRNAs and targets.

K-means clustering of miRNA expression showed 18 miRNAs present in a symbiosis downregulated cluster and 32 miRNAs in a ‘symbiosis upregulated’ cluster (Fig. 4b, Fig. S8b). We tested correlation in expression between each miRNA in these clusters and all of their predicted target genes. Out of 14,142 predicted target interactions, only 201 showed significant correlations in the relative expression of a miRNA and a predicted target gene (Pearsons, FDR <0.05). Of these, 82 correlations were negative, as expected for regulation via transcript cleavage. A subset of these miRNAs and predicted target transcripts with significant negative correlations are shown in Fig. 4c,d, and Fig. S9.

Twelve of sixteen MIR395 family miRNAs were in the symbiosis upregulated cluster, and nine showed significant negative correlations with at least one target gene (Fig. 4c). These putative miR395 targets broadly show connections to auxin biosynthesis and signaling. Among them is the nitrate transporter NRT1.1 (LotjaGi1g1v0065700), which in *Arabidopsis thaliania* has been shown to also negatively regulate auxin biosynthesis and polar transport in the root (Maghiaoui *et al*. 2020). MIR395 also targets STK (LotjaGi3g1v0010000) a putative kinase whose *At* ortholog (AtAGC1-3/PAX) is involved in auxin efflux regulation and root vascular development of protophloem via phosphorylation of PIN proteins (Marhava *et al*. 2018). An AGC kinase has also been shown to be specifically expressed in the proximal infection zone of *Medicago truncatula* nodules (Pislariu & Dickstein 2007) indicating a possible role for AGC kinases in regulating auxin distribution within nodules.

We identified eight mature miR2111 family miRNAs in the symbiotically downregulated cluster. Six had significant negative correlations with predicted target genes (Fig. 4d). The known role of lja-miR2111-5p is in the Autoregulation of Nodulation pathway as a repressor of TOO MUCH LOVE (TML) (Tsikou et al., 2018). We find a significant negative correlation between LjTML (LotjaGi1g1v0623800) and miR2111_4-5p. We also found a significant negative correlation between miR2111_7-3p and the MtSymCRK Cysteine Rich Receptor Like protein kinase homolog, which has been shown to be necessary for symbiosis in *Medicago trunculata* (Berrabah *et al*. 2014). This data suggests the possibility that the miR2111 family of miRNAs may have additional targets than previously known.

We identified two novel miRNAs which showed symbiosis-affected expression and had significant negative correlations with target genes. Novel_miRNA_17-5p is upregulated in symbiotic treatments and has a significant negative correlation with an ACT domain containing protein gene LotjaGi3g1v0281500_LC (Fig. S9a). Novel_miRNA_20-3p is downregulated in symbiosis treatments and is negatively correlated with LotjaGi5g1v0193900 (APG1), a putative methyltransferase, and LotjaGi1g1v0652300 (SEC14L-PITP) a putative phosphatidylinositol transfer family protein (Fig. S9b).

### CircRNAs show treatment specificity and differential expression

A total of 25,890 circRNAs were identified with 4,087 circRNAs consistently observed in at least three bioreplicates of the same treatment (Fig. S10, supplemental file 6). All individual treatments had more unique circRNAs than were shared between any set of treatments. The largets overlap was between G5R and S5N, followed by the set of circRNAs shared across all treatments (Fig. S11a). Among circRNAs detected in three bioreplicates of the same treatment, most circRNAs were exclusive to S5N (484) and G5R (184), with 79 consistently detected circRNAs shared across all treatments and 72 shared between G5R and S5N (Fig. 5a; Fig. S11b). This suggests that there is a population of circRNAs that are either widely detected, being candidates for ‘housekeeping’ circRNAs, and that S5N and G5R both show elevated numbers of consistent circRNAs compared to other treatments.

**Fig. 5.**
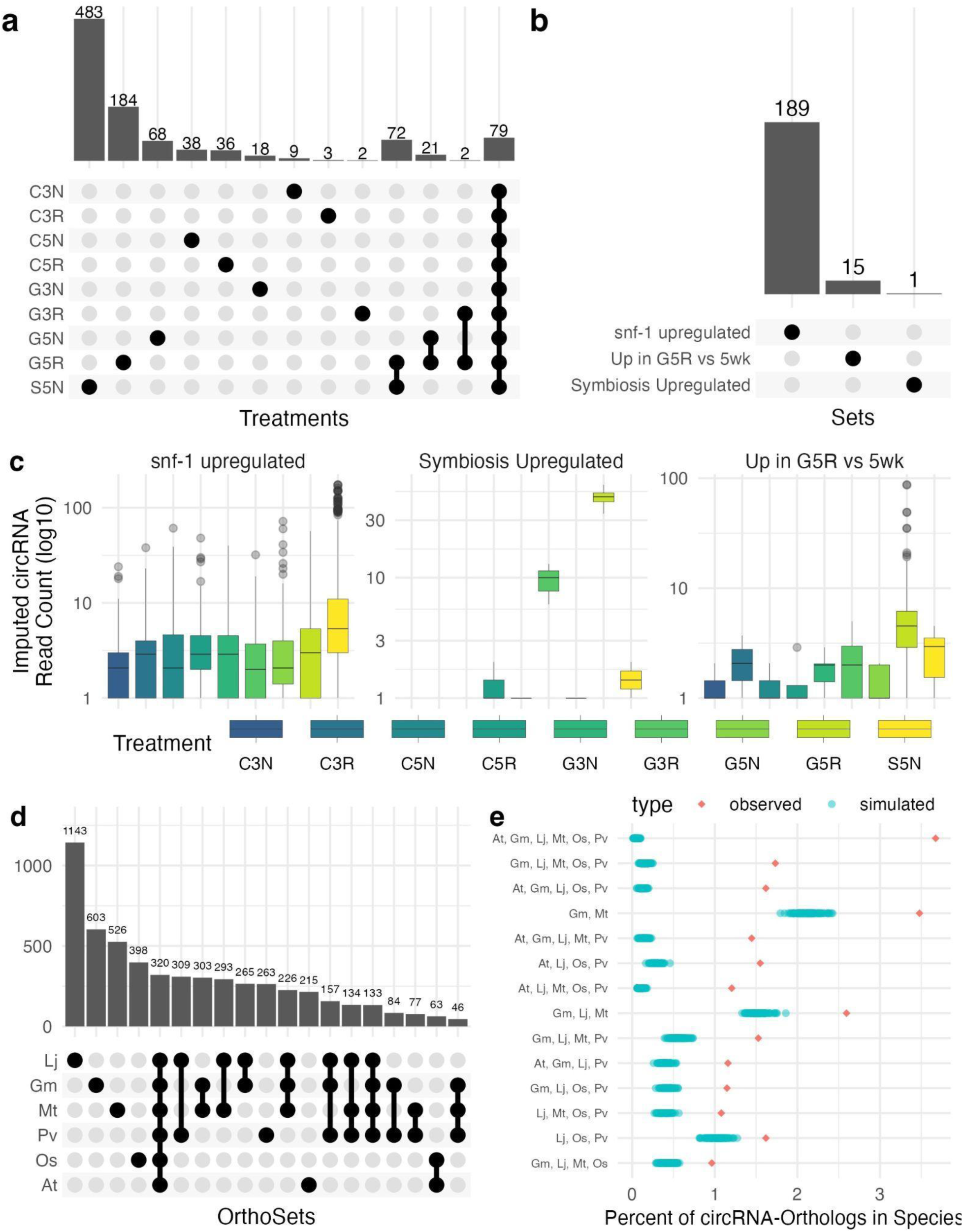
Differential expression of circRNAs. a) Consistently detected circRNAs across treatments. CircRNAs that are in the intersection of multiple treatments are not necessarily in three replicates of multiple treatments but were in at least three replicates of at least one treatment. b) Upset plot of the number of differentially expressed circRNAs in select treatment comparisons. c) Expression across treatments of the circRNAs that were upregulated in *snf-1*, upregulated in G5R versus other 5wpi treatments, or upregulated in symbiosis treatments. d) Intersections of orthogroup genes which act as circRNA parents in one or more species (At, *Arabidopsis thaliania;* Gm, *Glycine max;* Lj, *Lotus japonicus;* Mt, *Medicago truncatula*; Os, *Oryza sativa*; Pv, *Phaseolus vulgaris*), intersections shown here focus on comparisons between legumes, the 22 largest intersections are visible in Fig. S13a. e) Comparison of observed rates of shared circRNA-parent orthologs across species to simulated rates based on random assignment of circRNAs to genes for each species. Red diamonds to the right of blue points indicate that more conservation than is expected was observed. The 14 most divergent sets are shown here and the remaining intersection sets can be seen in Fig. S13b.

We identified 1,123 differentially expressed (DE) circRNAs (absolute log_2_(FC)≥1, BH Adj. P-value <0.05). Comparing DE circRNAs across treatments showed 189 *snf-1* upregulated circRNAs, 15 circRNAs upregulated in G5R versus all other 5wpi treatments, and a single circRNA which was upregulated in all symbiosis treatments compared to analogous treatments without symbiosis (Fig 4b,c).

### CircRNAs show conservation of parent genes

The degree of sequence conservation of circRNAs across plant species is difficult to establish due to a dearth of full-sequence circRNAs. An alternate measure of circRNA conservation is whether orthologous genes in different plant species produce circRNAs. We identified orthogroups for 8 plant species and obtained circRNA data for 5 plant species besides *Lotus japonicus*: *Arabidopsis thaliana, Glycine max, Medicago truncatula, Oryza sativa,* and *Phaseolus vulgaris*. Thirty-six percent of orthogroups were found in all 8 species with more conservation among angiosperms and legumes. Most orthogroups contained between 2-10 genes (Fig. S12a,b, supplemental file 7). Conservation of circRNA expression from orthologous genes was better-than-random. CircRNA-parent orthologous genes expressed a circRNA in all 6 species in 3.6% of cases compared to ∼0.05% in random simulations (Fig. 4d,e; Fig. S13a,b).

### *Lotus japonicus* circRNAs are likely predominantly *trans-*acting

To explore possible functions of circRNAs, we used their full-length sequences as input into psRNAtargetV2 to predict miRNA recognition elements (MREs). Among 14,115 unique circRNA isoforms 2,986 were predicted to have at least one MRE (Fig. S14a). Most circRNAs had no or only a few predicted MRE sites with one circRNA predicted to have 95 predicted MREs (Fig. S14b). In contrast, all 301 expressed mature miRNAs had at least one predicted circRNA target, with 14 miRNAs having more than 50 circRNA targets (Fig. S14c).

A linear RNA that is targeted by a miRNA may be stabilized by a circRNA that acts as a target mimic. We propose that this can happen in a *cis*- or *trans*-context. A gene with an expressed linear isoform targeted by a miRNA may also express a targeted circular RNA, which partially protects that linear isoform by acting as a target mimic. Alternatively, a *trans*-acting circRNA may be expressed from a different locus and still act as a target mimic for that miRNA. It is also possible that circRNAs expressed from genes have largely different MRE profiles than the linear isoforms due to differences in splicing. This is most obvious in the case of an MRE that overlaps or is very proximate to the back-splice junction; we have termed these MREs back-splice generated MREs (bsgMREs) (Fig. 6) (Utley *et al*. 2026). In theory, bsgMREs are expected to act in *trans* as the BSJ sequence motif is unique to the circRNA. In practice, the BSJ sequence may still exist somewhere within the linear cognate, and so bsgMREs may not be exclusive to *trans*-acting target mimicry. BsgMREs may also be additional sites for a miRNA that would still otherwise target the linear cognate, or they may be recognized by a miRNA that does not contain an MRE on the linear cognate and enable the circRNA to act as a trans-regulatory molecule (Fig. 6b).

**Fig. 6.**
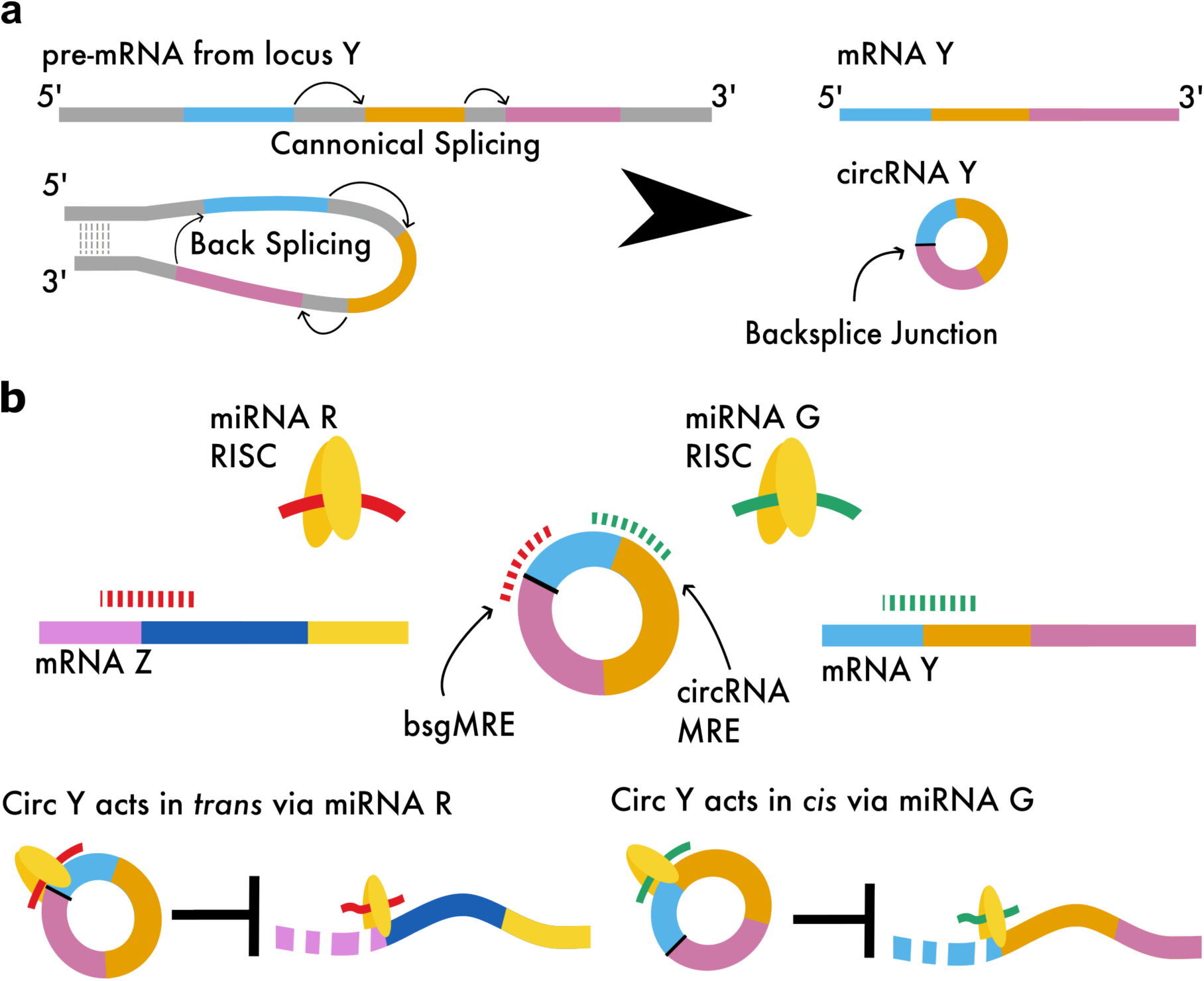
Diagram of circRNA cis and trans target mimicry. (a) Simplified diagram of splicing of pre-mRNA and back splicing of the same transcript to produce circRNA. (b) A diagram example of *cis-* and *trans-*acting circRNA MREs. The hypothetical circRNA circY is produced from the same genic locus as mRNA Y. CircY has a bsgMRE, which can be recognized by miRNA R, and an MRE recognized by miRNA G which is also shared by mRNA Y. Messenger RNA Z is expressed from a different gene and is targeted by miRNA R. If circRNA Y binds miRNA G and, in so doing, stabilizes mRNA Y, it is acting as a *cis-*regulatory target mimic. If circRNA Y binds miRNA R and, in so doing, stabilizes mRNA Z, it is acting as a *trans-*regulatory target mimic.

To identify whether *cis*- or *trans-*acting circRNAs are more predominant among bsgMREs in our dataset, we selected genes that expressed linear and circular RNAs with predicted MREs. For each miRNA-gene combination, we examined whether the miRNA targeted a linear isoform, a circular RNA through a normal MRE, and/or a circular RNA through a bsgMRE. Surprisingly, there was relatively little overlap in miRNA targeting between linear and circular RNAs expressed from the same gene. Out of 2,016 instances of miRNA-circRNA targeting per gene, only 408 (20%) had the same miRNA target a linear isoform from the same gene (Fig. 6a). This is surprising as it indicates that, at least for isoforms and circRNAs for which we have assembled sequences, the differences in splicing between circRNAs and linear RNAs leave relatively little MRE overlap. For bsgMREs there was even less co-targeting, out of 162 miRNAs that targeted at least one bsgMRE from a gene, only 9 also targeted a linear isoform from the same gene. These results indicate that circRNAs may function primarily as *trans*-acting regulatory molecules, at least via miRNA target mimicry.

### *Trans-*acting circRNAs are also more common in Arabidopsis thaliana *and* Oryza sativa

Is the predominance of *trans-*acting circRNAs similar in other plant species? We used publicly available circRNA and miRNA sequences from *Arabidopsis thaliana* and *Oryza sativa* and investigated circRNA:miRNA and miRNA:gene target predictions (Chu *et al*. 2022). Ratios for *cis*-acting and *trans*-acting circMREs and bsgMREs were found similar across the 3 species. All species have a majority of *trans-*acting circMREs which show no corresponding MRE for linear transcripts from the circRNA parent gene - 89% in arabidopsis, 91% in lotus, and 94% in rice (Fig. 6b). Again, bsgMREs showed a strong enrichment for *trans* activity, 97% of bsgMREs are *trans-*acting in *At*, *Lj*, and *Os*.

We also sought to identify whether specific circRNA:miRNA relationships may be conserved across species – similar miRNAs targeting circRNAs and linear transcripts from the same orthogroups across species. We clustered mature miRNAs by sequence from *At*, *Lj*, and *Os* to create comparable cross-species cluster ids. MiRNAs were fairly divergent and only 31 out of 813 miRNA clusters were present in all 3 species (Fig. S15a). MiRNA clusters predicted to target linear transcripts from the same orthogroup in all 3 species were relatively rare (103 instances out of ∼135,000), but include previously reported examples such as miR156 and miR535 targeting SPL genes and miR170/miR171 targeting GRAS family transcription factors (Fig. S15b., supplemental file 7) (Zheng, Ye, *et al*. 2019; Zhang *et al*. 2022; Pei *et al*. 2023).

Many miRNA clusters that were shared by all 3 species were predicted to target circRNAs or bsgMREs in all 3 species (29 out of 31, and 15 out of 31 respectively, Fig. S15c, d). However, shared targeting of orthologous circRNAs was very rare with only one common orthogroup with circRNA parents targeted by the same miRNA cluster in *At, Lj,* and *Os* (Fig. 7c). Among 2-species comparisons there were 4 targeting relationships in common between *At* and *Lj* circRNAs, 11 in common between *Lj* and *Os*, and 24 in common between *At* and *Os*. These conserved relationships are predominantly *trans*-acting. For example, miR394 putatively targets circRNAs from ADP-ribosylation factor genes in all 3 species and is predicted to target linear RNAs from Galactose oxidase/kelch repeat superfamily genes and loricrin-like protein genes in all 3 species, but is not predicted to target linear RNAs from ADP-ribosylation factor genes in *At*, *Lj*, or *Os*. Taken together these results indicate that *trans-acting* circRNAs may be predominant in plants and some *trans-acting* circRNAs may be part of conserved regulatory relationships, however both of these points require further analysis and validation.

**Fig. 7.**
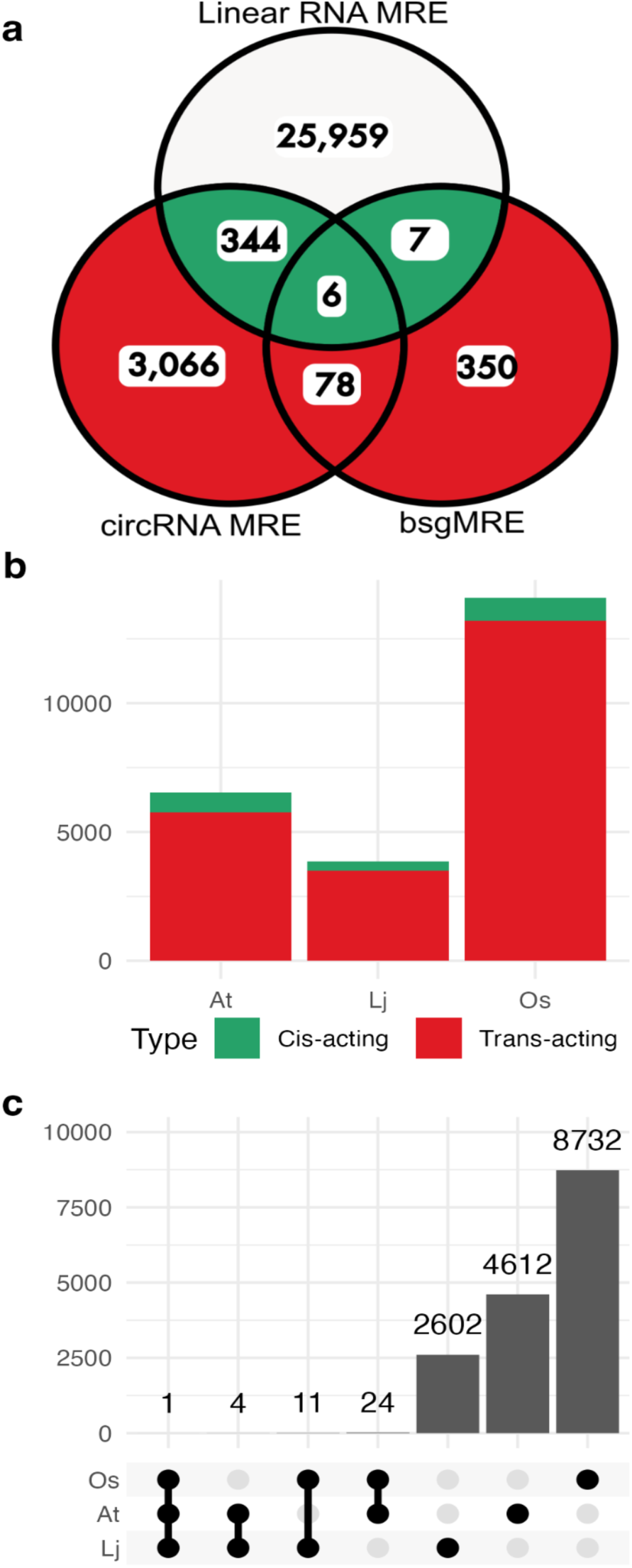
Comparisons of *cis-* and *trans-* acting circRNAs. a) Venn diagram displaying the number of MREs that target a given gene and/or circRNAs from that gene. Intersections colored in red show putatively *trans-*acting MREs that are distinct to a circRNA and do not have a corresponding MRE for the circRNA parent gene. Intersections colored in green indicate that these MREs are shared on both the circRNA and the circRNA parent gene. b) Counts of circRNA-MREs that are *cis-* or *trans*-acting from *Arabidopsis thaliana, Lotus japonicus,* and *Oryza sativa.* c) Intersections of miRNA clusters targeting conserved circRNAs.

### MREs of differentially expressed circRNAs are predominantly *trans-*acting

*Trans-*acting circRNAs are potentially interesting regulatory molecules and their differential expression may offer insight into their regulatory functions. Among the 15 DE circRNAs in the “Up in G5R vs 5wk” group, only two were predicted to contain MREs, both predicted to be *trans-acting*. Of the 181 *snf-1* upregulated circRNAs 81 contained at least one MRE and 5 contained a bsgMRE (Supplemental Table 1). Only two MREs on these 81 circRNAs were predicted to be *cis-*regulatory, both located on the same *snf-1* upregulated circRNA (LjG1.1_chr1:88664207-88665094; LotjaGi1g1v0399500), including one predicted bsgMRE for novel_miRNA_35-5p.

### CircRNA Validation

We chose to validate a selection of circRNAs identified by deep sequencing using divergent RT-PCR based on evidence of differential expression and presence of predicted MREs and bsgMREs. Among these were one symbiosis upregulated circRNA, 2 circRNAs that were upregulated in the symbiosis treatments at 5wpi with predicted MREs, and 2 *snf-1* upregulated circRNAs with predicted bsgMREs.

Our standard of evidence for validating these circRNAs is to recover amplicons containing the backsplice-junction from divergent primer RT-PCR and verify the same primers did not produce the same amplicon from genomic DNA. The symbiosis-upregulated circRNA is expressed from a putative potassium transporter (LotjaGi4g1v0016300) the full sequence was validated as all of exon 2 and is designated as circKT1(2) (Fig. S17a). One of the circRNAs upregulated in the G5R treatment compared to other 5wpi treatments is expressed from a putative serine threonine kinase gene (LotjaGi1g1v0054700) and is composed of two 5’ UTR regions and exons 1-3 and is designated circSTK(5U1,5U2,1,2,3) we also validated a circRNA which lacks one of the 5’ UTR regions, circSTK(5U2,1,2,3) (Fig. S17b). We validated a *snf-1* upregulated circRNA expressed from a putative Mediator of RNA Polymerase II subunit 13 gene (LotjaGi6g1v0031200) circMED13(2,3,4,5,6) and confirmed the sequence for predicted bsgMREs for miR395_2-5p and novel_miRNA_29-3p (Fig. S17c). We attempted to validate another *snf-1* upregulated circRNA from a putative glycolipid transfer gene (LotjaGi5g1v0189300). We were unable to recover the exact circRNA identified via CIRI-full. However, we validated 2 similar circRNAs with the same 3’ side of the back-splice junction, one of which maintains a miR5574 bsgMRE identified in the original CIRI-full circRNA (Fig S17d).

Another circRNA upregulated in symbiotically active treatments at 5wpi was a + strand circRNA overlapping in the opposite orientation of *LjnsRING*/BRUTUS (LotjaGi4g1v0453800). We were also interested in the strandedness of linear RNAs in the region and examined the locus in 4 windows ranging from the 3’ end of an upstream + strand gene (LotjaGi4g1v0453700) through to exon 7 of BRUTUS. In each of these windows we performed RT with single + and - primers and performed PCR with convergent primers for all windows as well as divergent primers for exon 11 to validate the circRNA (Fig. 7a). This provides evidence for a BRUTUS-antisense + strand RNA that extends from exon 14 to at least exon 7 and does not include the BRUTUS 3’ UTR. The divergent RT-PCR reactions revealed identical products in both + and - strand reactions that were present in both RNAseR treated and untreated reactions. This validates the expected + strand circular RNA circBRUT(10,11;+) as well as a larger circRNA, circBRUT(10,11,12,13;+) and the - strand equivalents; circBRUT(10,11;-) and circBRUT(10,11,12,13;-) (Fig. 7b,c). Genomic DNA PCR for all convergent primers produced the expected results which in total shows that this locus is in agreement with the reference. Divergent primer gDNA PCR produced amplicons that were largely PCR artefacts - containing flanking intronic regions with self-complementarity. However, one gDNA PCR band was the same size as circBRUT(10,11) and was a similar sequence including the bsj region and lacking introns; sanger sequencing from this band showed multiple peaks and separating the peaks showed a portion of sequence that contained introns. Interpreting this gDNA product is difficult, but it could only be conflated with one of the two detected circRNA compositions. Because the locus itself is free from tandem genomic duplications and the divergent RT-PCR products were present in DNAse and RNAseR treated samples we consider the circRNAs from BRUTUS to be well supported.

### Competing endogenous RNA network shows circRNAs with potential regulatory functions

CircRNAs and non-coding RNAs may act as target mimics in competing endogenous RNA (ceRNA) networks to regulate genes. To examine the specific potential functions of our ncRNAs in symbiosis we constructed ceRNA network models based on predicted MREs on linear and circular RNAs. The total predicted network was composed of 29,394 nodes and 105,621 edges (supplemental file 7). To focus on the portion of the network potentially relevant to symbiosis we limited the network to only those genes, circRNAs, and ncRNAs which showed symbiosis-specific expression and their associated miRNAs. This reduced the network to 905 nodes and 1190 edges (Fig. 8a). From this filtered network we examined two local subnetworks around circRNAs and ncRNAs (Fig. 9b, c).

**Fig. 8.**
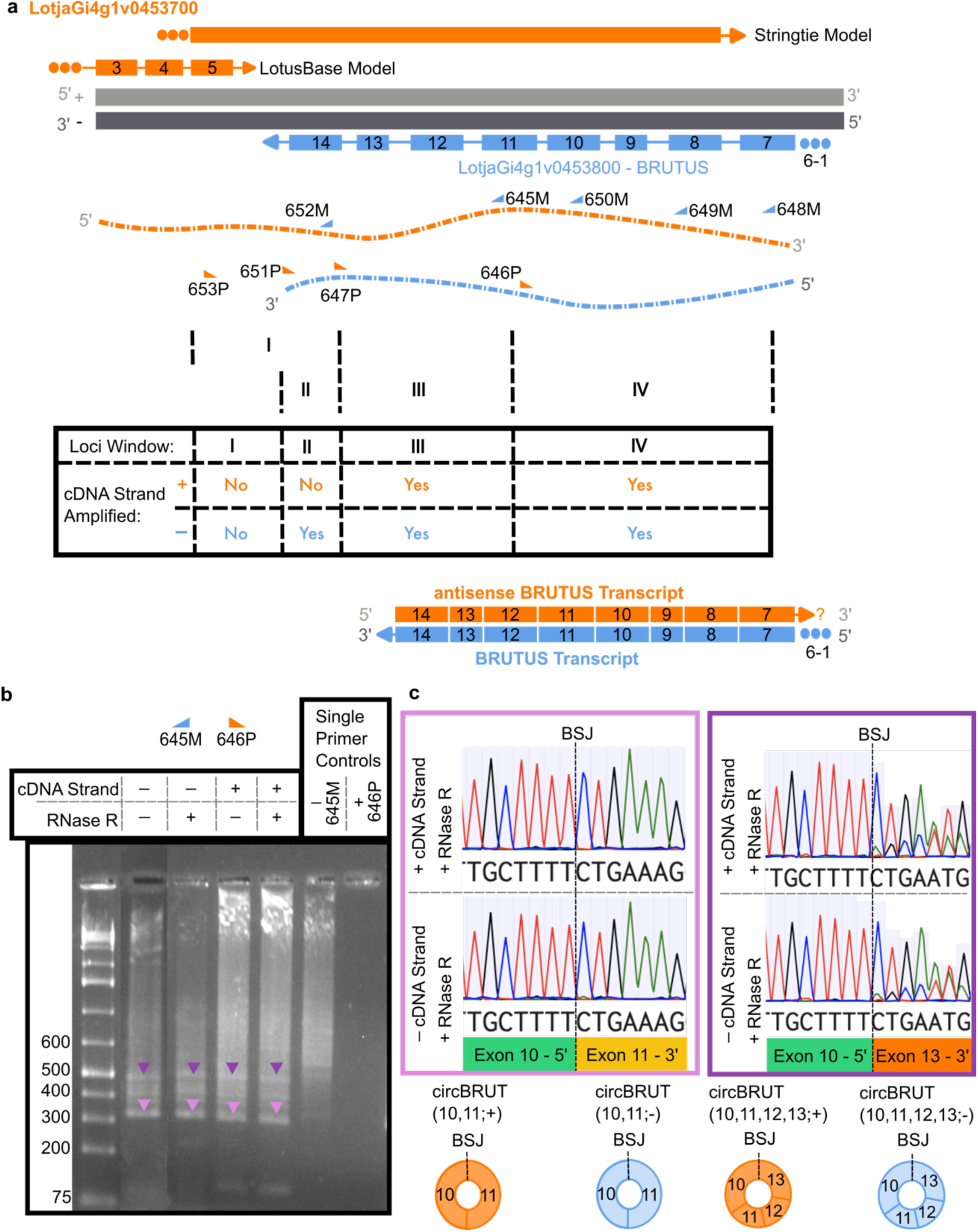
BRUTUS linear and circular RNA. (a) Diagram of BRUTUS (LotjaGi4g1v0453800) locus transcript models. LotjaGi4g1v0453700 is encoded on the + strand and is depicted in orange with two different transcript models: LotusBase and Stringtie assembled transcripts generated from our RNAseq data. The Stringtie transcript model has a tail-to-tail overlap with the BRUTUS transcript. The BRUTUS transcript model is shown in blue and is encoded on the - strand. Diagrams of RNA strands from the + and - strands corresponding to the annotated genes are shown along with primer names and locations. Primers ending in P are + strand primers and primers ending in M are - strand primers. The locus was divided into 4 regions (I-IV) which were examined with single-primer first-strand cDNA synthesis. The results of these RT-PCR reactions are summarized in the embedded table. This resulted in the discovery of a BRUTUS anti-sense (+-strand) RNA that runs from exon 14 to at least exon 7 of BRUTUS (Fig. S19). (b) Gel showing divergent RT-PCR for divergent primers for + and - strand cDNA PCR with and without RNAse R treatment. Purple and pink triangles indicate bands that were Sanger sequenced. (c) Excerpts of Sanger sequence alignments which cover the BSJs and circRNA diagrams for circBRUT(10,11) and circBRUT(10,11,12,13).

**Fig. 9.**
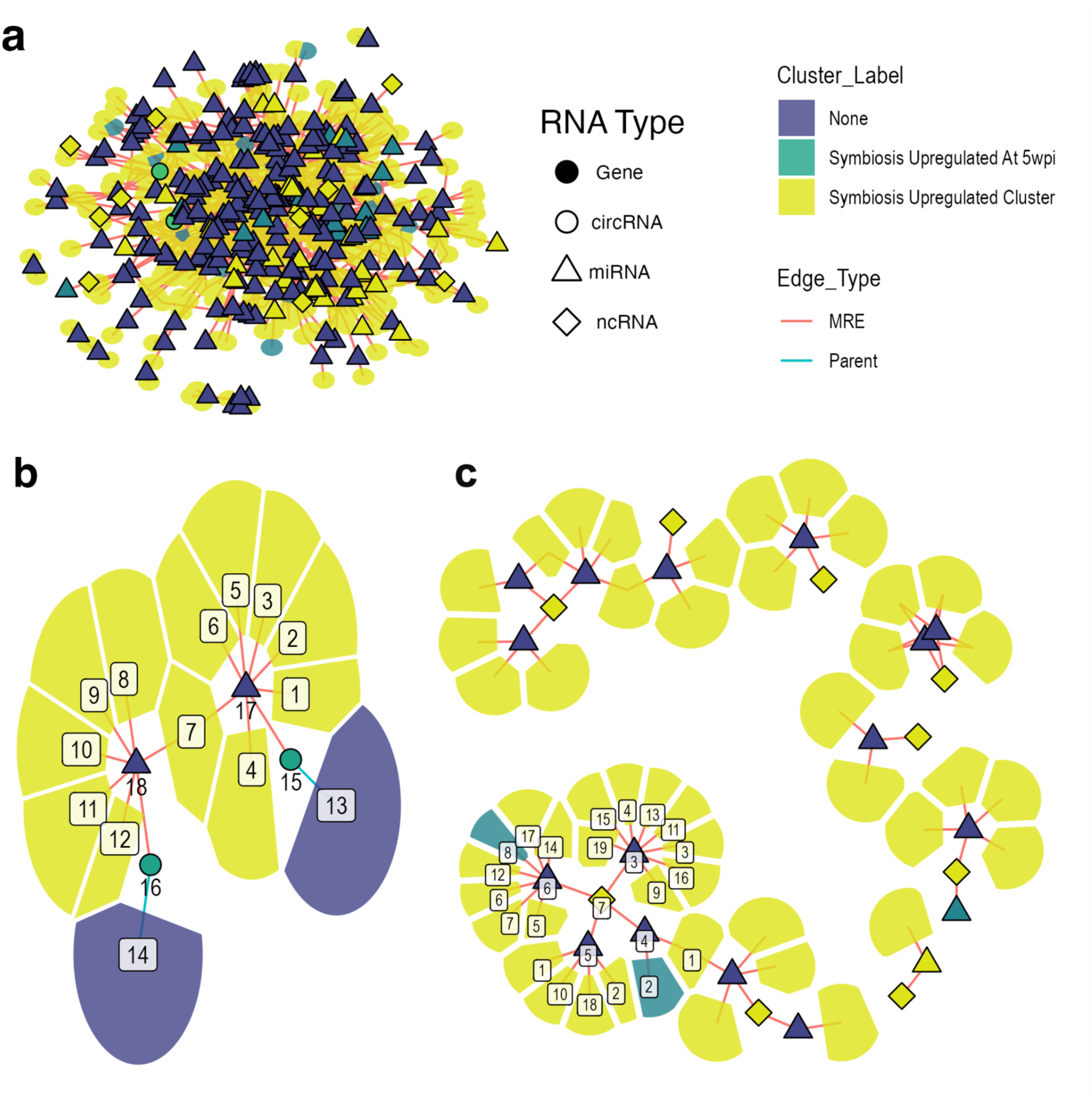
Competing endogenous RNA Network Model. (a) Network of genes, circRNAs, and non-coding RNAs which are predicted targets of miRNAs and belong to symbiotically up- or downregulated clusters. B and C are local subnetworks of A. B) Shows the local network (up to two connecting edges) around the only two circRNAs which are upregulated in G5R with predicted MREs. CircRNA parent genes are also shown in the network and are not targeted by either miRNA in the local network indicating that these circRNAs are purely trans-regulatory in terms of miRNA binding activity. C) shows the local network (within 2 connecting edges) around all 9 non-coding RNAs which are in the symbiotically upregulated expression cluster and have MREs. All nodes in B and select nodes in C are given an index number and further information is available in supplemental file 7.

The circRNA local network includes validated circRNAs expressed from a BRUTUS homolog (circBRUT(10,11;+)) and expressed from a putative serine threonine kinase (circSTK(5U1,5U2,1,2,3), LotjaGi1g1v0054700). Both circRNAs are upregulated under symbiosis treatment at 5 weeks post-inoculation. Neither parent gene is expected to be targeted by the miRNAs that are predicted to target the circRNAs, indicating that both circRNAs are potentially trans-regulatory. Novel_miR_9-3p is predicted to target circBRUT(10,11;+) and 6 symbiotically upregulated genes. The miRNA miR3623_1-3p is predicted to target circSTK(5U1,5U2,1,2,3) and 7 symbiotically-upregulated genes. Both novel_miR_9-3p and miR3623_1-3p are predicted to target LjPMT1, a putative polyol transporter. It is possible that circBRUT(10,11;+) and circSTK(5U1,5U2,1,2,3) participate in the regulation of LjPMT1 by sequestering novel_miR_9-3p and miR3623_1-3p which do not show symbiosis specific expression.

We identified 9 symbiotically upregulated ncRNAs with predicted MREs. LotjaGi1v1ncRNA14 is a 4kb non-coding RNA that has 4 predicted MREs. All of the miRNAs predicted to target LotjaGi1v1ncRNA14 also target symbiosis upregulated genes. Among these symbiosis upregulated targets is LotjaGi6g1v0341200 a putative equilibrative nucleoside transporter whose transcripts were specific to the infected zone of the nodule (Ye *et al*. 2024). Nucleosides can be catabolized as a nitrogen source or used as a source of nucleobases in either the plant or the bacteroid. Nucleoside transport may also be part of signaling or cytokinin regulation (Korobova, A *et al*. 2021; Dunken *et al*. 2024). There are also several symbiosis upregulated co-targets of LotjaGi1v1ncRNA14 that may have roles in regulating transcription and splicing including LotjaGi2g1v0231600 (putative EFFECTOR OF TRANSCRIPTION2 ortholog), LotjaGi1g1v0091400 (AAR2 family protein), LotjaGi0g1v0000800 (Agenet domain protein). This network approach allows us to nominate circBRUT(10,11;+), circSTK(5U1,5U2,1,2,3), and LotjaGi1v1ncRNA14 as potential SNF-associated gene regulatory molecules.

## DISCUSSION

Various classes of non-coding RNAs are regulatory molecules in plants (Yadav *et al*. 2024). We sought to identify and computationally delineate non-coding RNAs that may have a potential regulatory role in the symbiosis between *Lotus japonicus* and *M. loti*. This work reveals 64 putatively novel miRNAs and shows differential expression of known and novel miRNAs in response to symbiosis. To predict MREs we used psRNAtargetV2 which has been shown to have good recall of known interactions but may be lacking in precision (Dai *et al*. 2018). We consider these predicted interactions as grounds for hypothesis formation, which require validation.

Differentially expressed miRNAs with negative correlations to predicted target genes add confidence to these predictions. Nine miR395 family microRNAs were upregulated under symbiosis and had negative correlations with predicted targets involved in auxin regulation including NRT1.1, STK, CYP711A1 (AtMAX1), and FMO (AtYUC6) suggesting a potential role for miR395 regulation of auxin signaling to support symbiosis (Lazar & Goodman 2006; Pislariu & Dickstein 2007; Marhava *et al*. 2018; Maghiaoui *et al*. 2020; Dai *et al*.). Regulation of auxin is involved in infection and nodule development and further studies could clarify if and how miR395 affects auxin distribution during these processes (Nagata & Suzuki 2014; Breakspear *et al*. 2014).

MicroRNAs with negative correlations to predicted target genes may also be involved in regulating membranes to support symbiosis. Novel_miRNA_20-3p is downregulated in symbiosis treatments and is negatively correlated with LotjaGi1g1v0652300 (SEC14L-PITP) a putative phosphatidylinositol transfer family protein. We observed a similar pattern of expression in miR156d_1-3p and the predicted target VAMP4 (LotjaGi3g1v0459500) (Fig. S9c). SEC14L-PITPs are membrane bound proteins which recognize and transport phospholipids and participate in membrane identity (Montag *et al*. 2023). Some SEC14L-PITP genes show nodule specific expression in *Lotus japonicus* (Kapranov *et al*. 2001). VAMP proteins are essential for symbiosome membrane formation in *Medicago trunculata* (Ivanov *et al*. 2012). Further studies could knockout SEC14L-PITP and VAMP4 or overexpress novel_miRNA_20-3p and miR156d_1-3p to examine their involvement in membrane formation or maintenance within nodules.

Circular RNAs have previously been shown to participate in competing endogenous RNA networks in plants (Zhou *et al*. 2021; Yin *et al*. 2024). We expand this concept by introducing the idea of *cis-* and *trans-*regulatory circRNA MREs. *Cis-*regulatory as we define it means that the same miRNA is predicted to target circular and linear transcripts originating from the same gene. *Trans-*regulatory circRNA MREs exist on the circRNA but there are no corresponding MREs on linear transcripts from the parent gene. Because circRNAs are derived from linear transcripts of a gene we expected that *cis-*regulatory MREs would be predominant, except for bsgMREs. Surprisingly, we found that the majority of predicted circRNA MREs were *trans-*regulatory across *Arabidopsis thaliana, Lotus japonicus,* and *Oryza sativa*. Predicted miRNA:circRNA relationships which are conserved in multiple species are also often *trans*-regulatory. Analysis of bsgMREs revealed additional *trans*-regulatory circRNAs. *Trans-*regulatory circRNAs may be a method for fine-tuning the expression of networked genes via the expression of and circRNA biogenesis from the circRNA parent gene. Our analysis suggests some circRNAs have conserved *trans-*regulatory roles across plant species. Validation and further analysis could help prioritize circRNAs which may be conserved functional molecules and useful candidates for genetic engineering.

Two potential *trans-*regulatory circRNAs are upregulated in symbiotically active treatments compared to analogous treatments and may act as target mimics for LjPMT1 through novel_miR_9-3p and miR3623_1-3p. LjPMT1 (LotjaGi4g1v0116000) was identified as expressed in the nodule cortex, infection zone, and the nitrogen fixation zone and is orthologous to the *Lj* MG20 ecotype LjPLT11 (Lj4g3v0937820) which was found to be highly expressed in nodule tissue (Tian *et al*. 2017; Ye *et al*. 2024). LjPLT11 is a peribacteroid membrane-localized d-pinitol transporter which contributes to nodule longevity (Tian *et al*. 2022).

One of the validated circRNAs that is a co-target of LjPMT1 was circBRUTUS(10,11;+). In lotus LjnsRING/BRUTUS is necessary for successful nodulation (Shimomura *et al*. 2006). The *Glycine max* ortholog BTSa has been shown to act as an iron sensor which positively regulates nodulation by stabilizing NSP1a proteins under iron-sufficient conditions (Ren *et al*. 2025). We identified sense and anti-sense circular RNAs expressed from the BRUTUS locus as well as a longer anti-sense RNA that is likely linear. The BRUTUS locus may therefore have both protein-coding and non-coding transcripts that regulate symbiosis. Knockdown and transgenic partitioning of the coding and non-coding transcripts of BRUTUS could be used to test this hypothesis. If circBRUT(10,11;+) acts as a target mimic it is expected that knockdown of circBRUT(10,11;+) would destabilize LjPMT1 transcripts and lead to early senescence of nodules.

To our knowledge this is the first report of otherwise identical + and - strand circRNAs. Similar circular RNA molecules of both polarities exist in plants as retrozymes, however the BRUTUS locus lacks the characteristic hammer-head ribozyme motifs of retrozymes (Cervera *et al*. 2016). Our methods do not show whether these circRNAs occur in the same cells or form a duplex, leaving open questions about localization, function, and biogenesis. Determining this with enzymatic or analytic methods would likely require native purification of the circRNAs and probe-based purifications may be hampered if a native duplex is present (Gabryelska *et al*. 2024). Alternatively, measuring the expression of both strands of circular and linear RNAs from the BRUTUS locus with cellular resolution may give a clear answer as to whether the + and - strand transcripts co-occur. While single cell methods have been devised that capture circular RNAs they have not yet been applied in plants (Fan *et al*. 2015; Xu *et al*. 2024).

## Supporting information

SF2 - Primers and PCR conditions

SF3 - M. loti transcript information

SF4 - Gene and Isoform information

SF5 - miRNA information

SF6 - CircRNA information

SF7 - Orthogroups, and miRNA predicted targeting information

## Abbreviations

(AON): Autoregulation of Nodulation
(bsgMRE): backsplice-generated miRNA response element
(circRNA): circular RNA
(ceRNA): competing endogenous RNA
(DEG): Differentially Expressed Genes
(GO): Gene Ontology
(MRE): miRNA response element
(lncRNAs): long non-coding RNAs

## SUPPLEMENTARY MATERIALS

**S1. Supplementary Methods**

**S2. Primers and PCR conditions;** S2.1 Primer Sequences; S2.2 BRUTUS locus Strand Specific RT-PCR information; S2.3 CircRNA RT-PCR information for other loci.

**S3. *M. Loti* transcript information**; S3.1 CPM *M. loti* genes; S3.2 LogFC *M. loti* genes; S3.3 *M. loti* DEG upset groups; S3.4 *M. loti* symbiosis DEG GO term enrichment.

**S4. Gene and isoform expression and annotations;** S4.1 Gene Annotations; S4.2 Gene Counts; S4.3 Filtered gene log2cpm; S4.4 Clustered DEGs; S4.5 Fig3b gene IDs S4.6 Isoform Peptide level annotations; S4.7 Isoform counts; S4.8 Filtered isoform log2cpm; S4.9 Clustered DE isoforms; S4.10 Comparison of DEG and DE isoform clusters.

**S5. miRNA Information;** S5.1 miRNA loci; S5.2 miRNA counts per million; S5.3 miRNA log2cpm and k-means clustering; S5.4 miRNA pairwise logFC; S5.5 miRNA DE upset information.

**S6. Observation level circRNA information;** S6.1 All CircRNA observations; S6.2 Imputed counts of circRNAs; S6.3 Imputed counts of circRNAs Log2(CPM); S6.4 Differentially Expressed circRNA Upset Groups.

**S7. Orthogroups, miRNA cluster orthogroup targeting, and ceRNA networks;** S7.1 Orthogroups; S7.2 At, Os, and Lj miRNA clusters; S7.3 miRNA cluster orthogroup targeting; S7.4 Fig. 9A Symbiosis Regulated Subnetwork; S7.5 Fig. 9B Symbiosis Regulated circRNA local network; S7.6 Fig. 9c Symbiosis Regulated lncRNA local network.

