## Supplemental Figures for "Identification of Potential Regulatory Non-Coding RNAs in *Lotus Japonicus* Symbiosis"

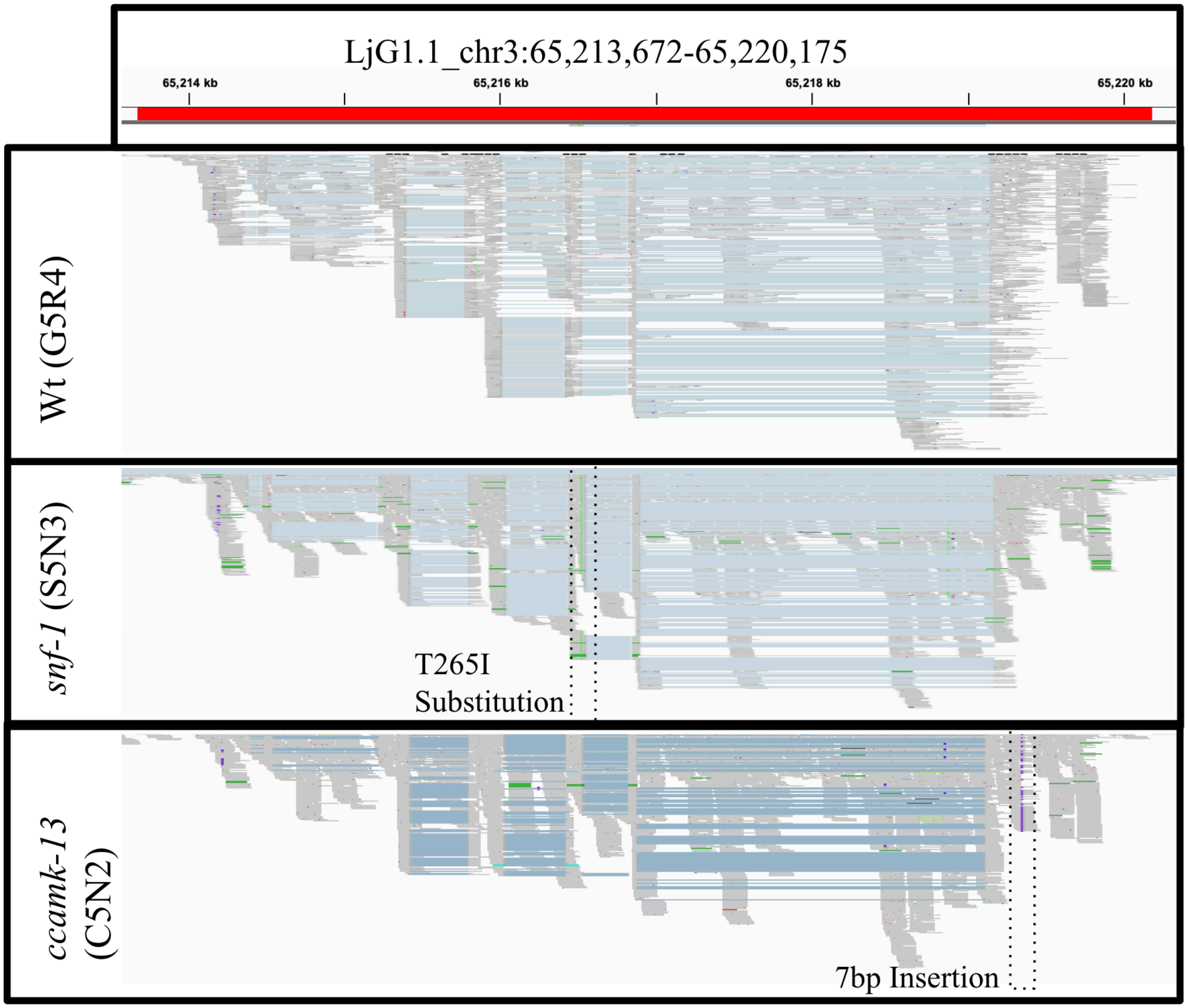


**Fig. S1.** RNAseq alignments to LjCCaMK (*LotjaGi3g1v0307700*) locus confirms mutant sequences and gene expression. RNAseq alignments from Wt, *snf-1*, and *ccamk-13* genotypes were inspected at the *LotjaGi3g1v0307700* locus and showed the expected mutations. The *snf-1* mutant shows the T265I substitution which is visible in the alignment as a vertical green band within the portion highlighted by dotted lines. The *ccamk-13* mutant shows the 7bp insertion which is visible as a vertical purple band within the highlighted portion.

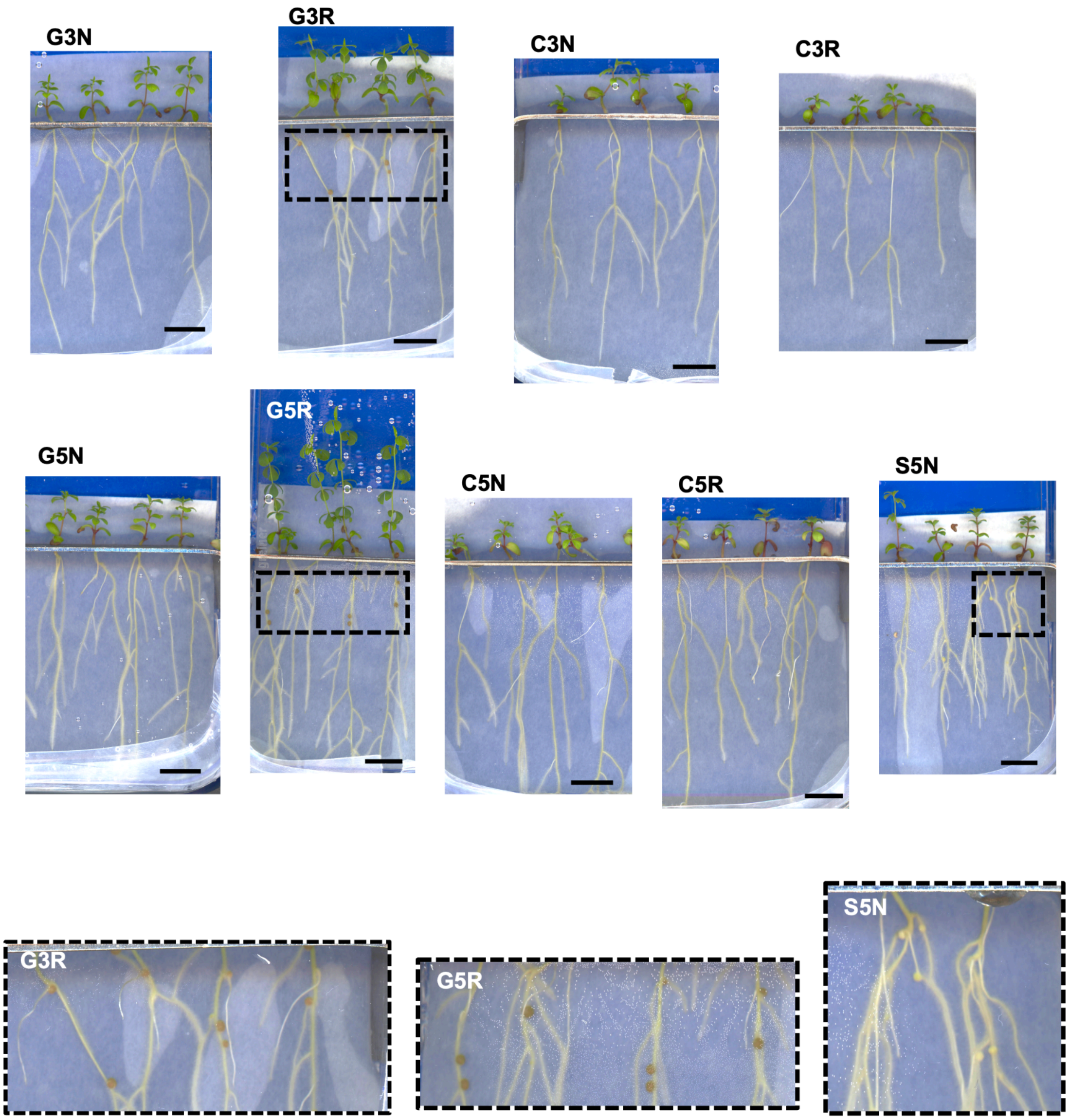


**Fig. S2.** Representative images of Lotus japonicus plants shortly before harvest. Images are labeled with the treatment condition, bar indicates 1cm. Outset images are zoomed to show nodules and nodule color. White nodules indicate that hemoglobin is absent and there isn't active symbiotic nitrogen fixation.

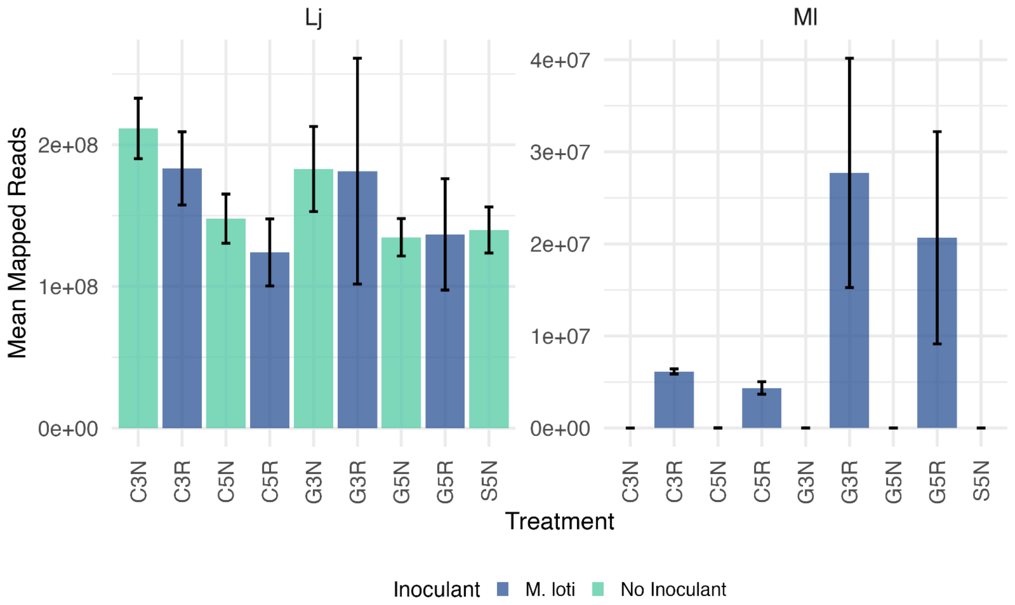


**Fig. S3.** Number of reads mapped to *Lotus japonicus* and *Mesorhizobium loti* from BBsplit. *Lotus japonicus* (Lj) is displayed at left and *Mesorhizobium loti* at left. The number of mapped reads was averaged per treatment, bars show 1 standard deviation. Bars are colored by the inoculant supplied to that treatment.

a

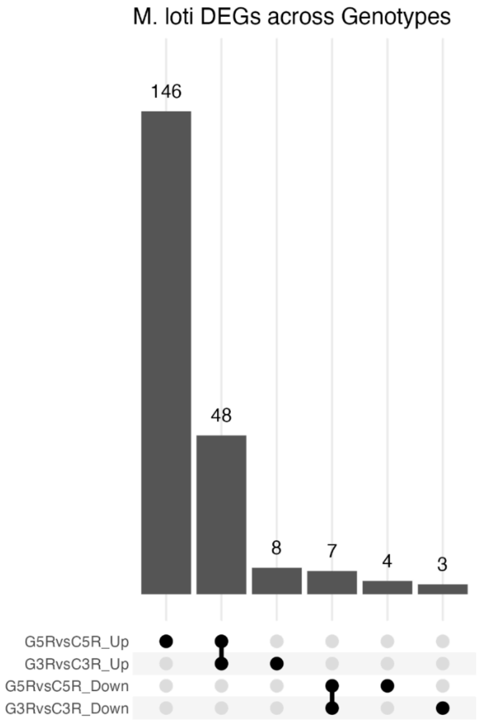


b

| Gene | Function | G3R vs C3R | G5R vs C5R | G5R vs G3R | C5R vs C3R | context |
|---|---|---|---|---|---|---|
| FlgE | flagellar hook proteins | -0.3 | -0.02 | 0.5 | 0.4 | Free living mobility |
| FlgK | flagellar hook associated proteins | -0.3 | -0.02 | 0.6 | 0.4 | |
| | flagellin | -0.2 | 0.0 | 0.9 | 0.8 | |
| NirD | nitrate ABC transporter permease | -1.8 | -0.8 | 2.2 | 1.2 | Assimilated N transport |
| | nitrite reductase small subunit | -1.6 | -1.3 | 1.3 | 1.0 | |
| NifE | nitrogenase Fe-Mo cofactor biosynthesis | 2.4 | 4.3 | 1.9 | 0.0 | Bacteroid N2 fixation |
| | nitrogenase Mo-Fe protein alpha chain | 2.4 | 5.0 | 2.4 | -0.3 | |
| | nitrogenase Mo-Fe protein beta chain | 3.0 | 4.9 | 1.9 | -0.1 | |
| Ccb3 | cytochrome-c oxidase, ccb3-type SUI | 2.0 | 4.1 | 1.4 | -0.7 | Respiration |
| | efflux RND transporter permease | -1.6 | -1.7 | 0.8 | 0.9 | Nod factor |
| | tRNA-Met | 2.5 | 1.4 | -0.2 | 0.9 | Translation |
| | tRNA-Gly and 18 other tRNAs | 1.7 | 1.5 | -0.4 | -0.2 | |
| | 50S ribosomal protein L5 and others | -0.8 | 0.3 | 2.4 | 1.3 | Translation |
| | 30S ribosomal protein and S10 and others | -0.9 | 0.1 | 2.3 | 1.3 | |

**Fig. S4.** Expression of *M. loti* genes. (A) Upset plot showing the number and intersections of differentially expressed *M. loti* genes across treatment comparisons. Labels indicate the treatment comparison and direction of gene expression (Ex. G5RvsC5R_Up indicates the number of genes that were upregulated in G5R compared to C5R). (B) Table of select differentially expressed *M. loti* genes relating to free-living or nitrogen-fixing functions. The terminal oxidase of the respiratory chain, *ccb3*, which has a higher affinity for oxygen than other terminal oxidases due to the need to bind $O_2$ as an electron acceptor in the low-oxygen environment of the leghemoglobin-containing bacteroids, was several-fold more highly expressed under inoculated Gifu treatments than under *ccamk-13* treatments.

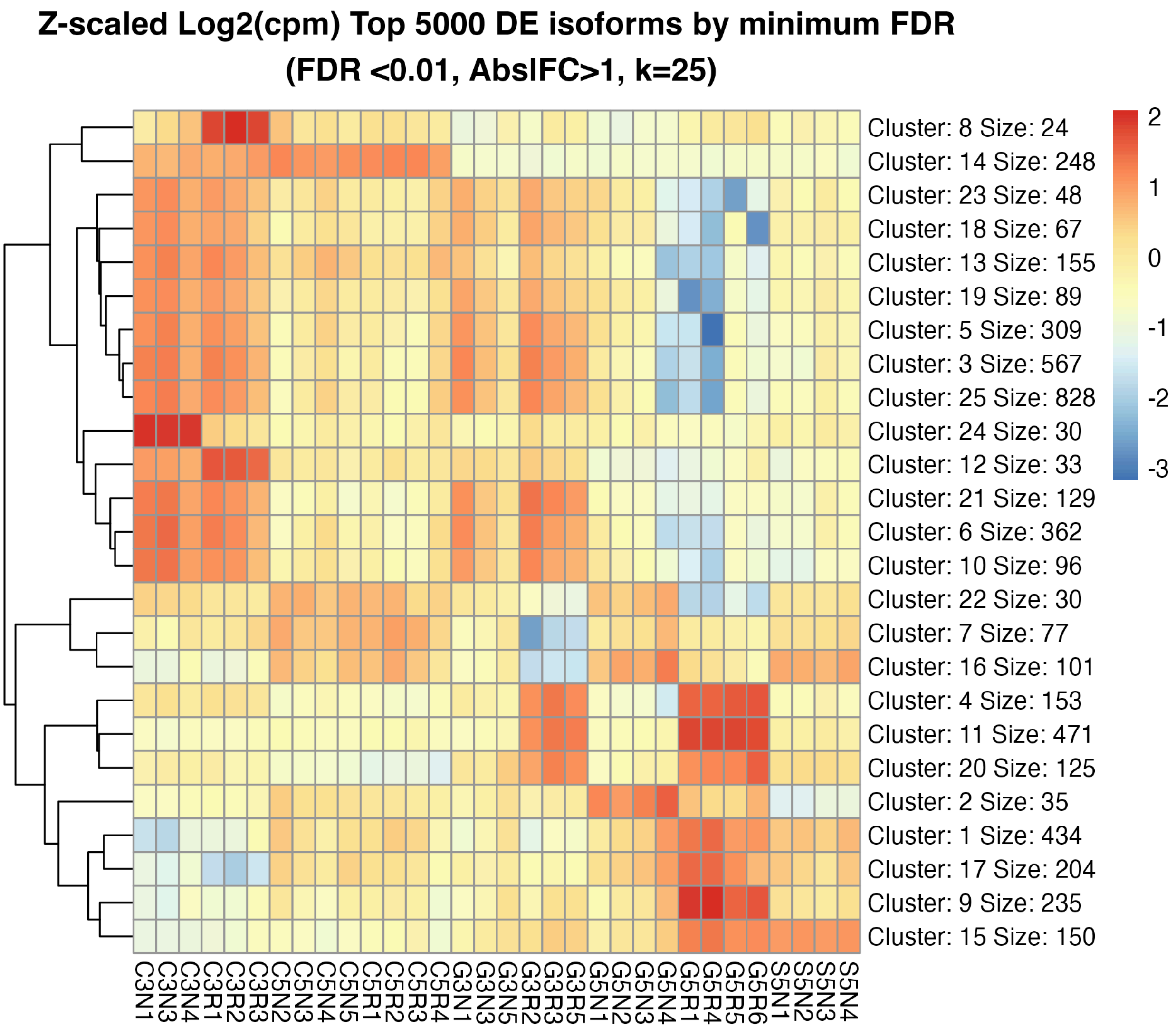


**Fig. S5.** K-Means clustering of the Top 5000 differentially expressed transcripts by Minimum FDR. Color shows relative expression as Z-scaled $Log_2$(Counts Per Million Mapped Reads).

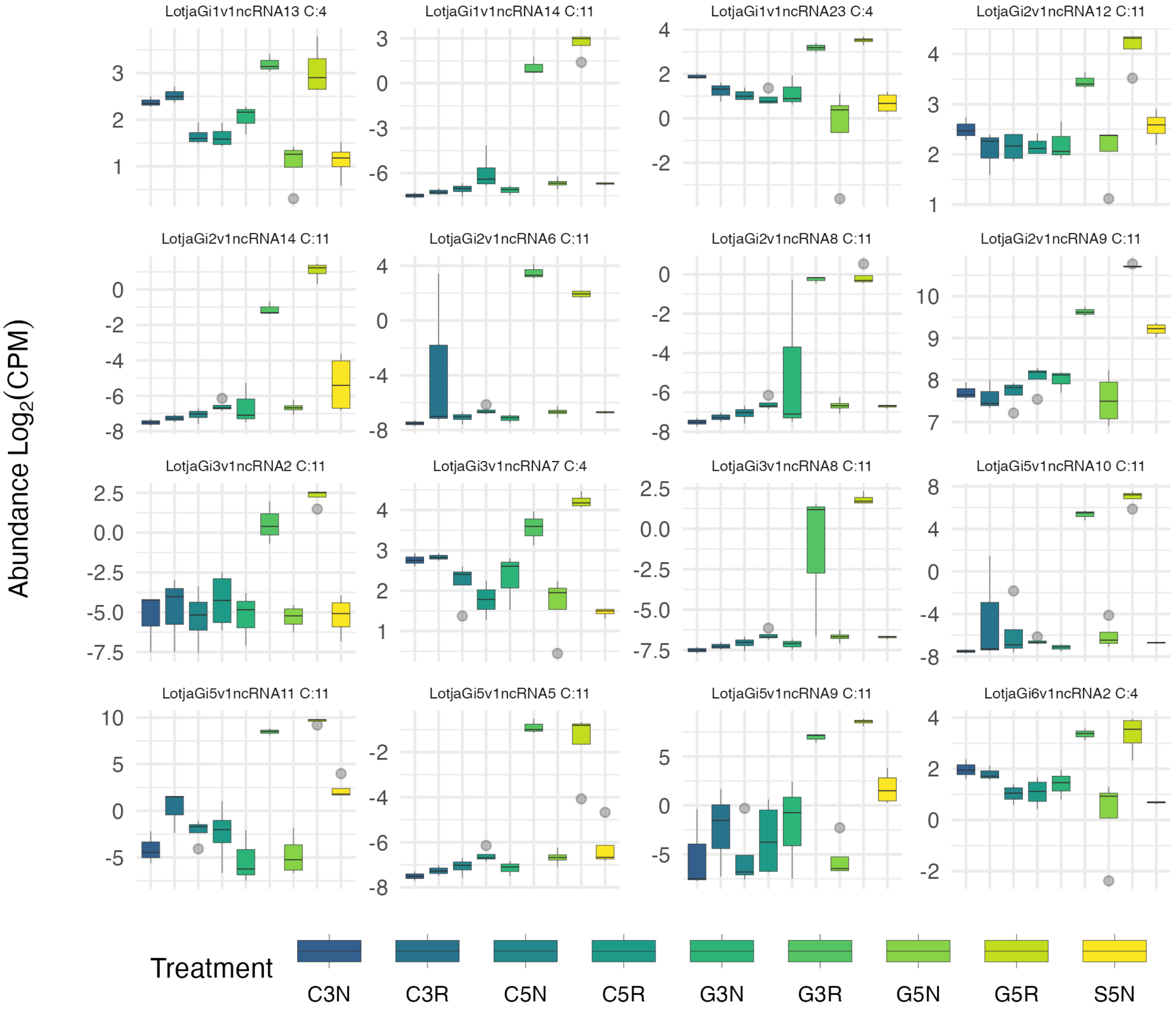


**Fig. S6.** Boxplots of expression of symbiosis-upregulated non-coding RNAs. Abundance is shown as $Log_2$(Counts Per Million Mapped Reads). N=3 for C3N, C3R, G3N, G3R and N=4 for all other treatments.

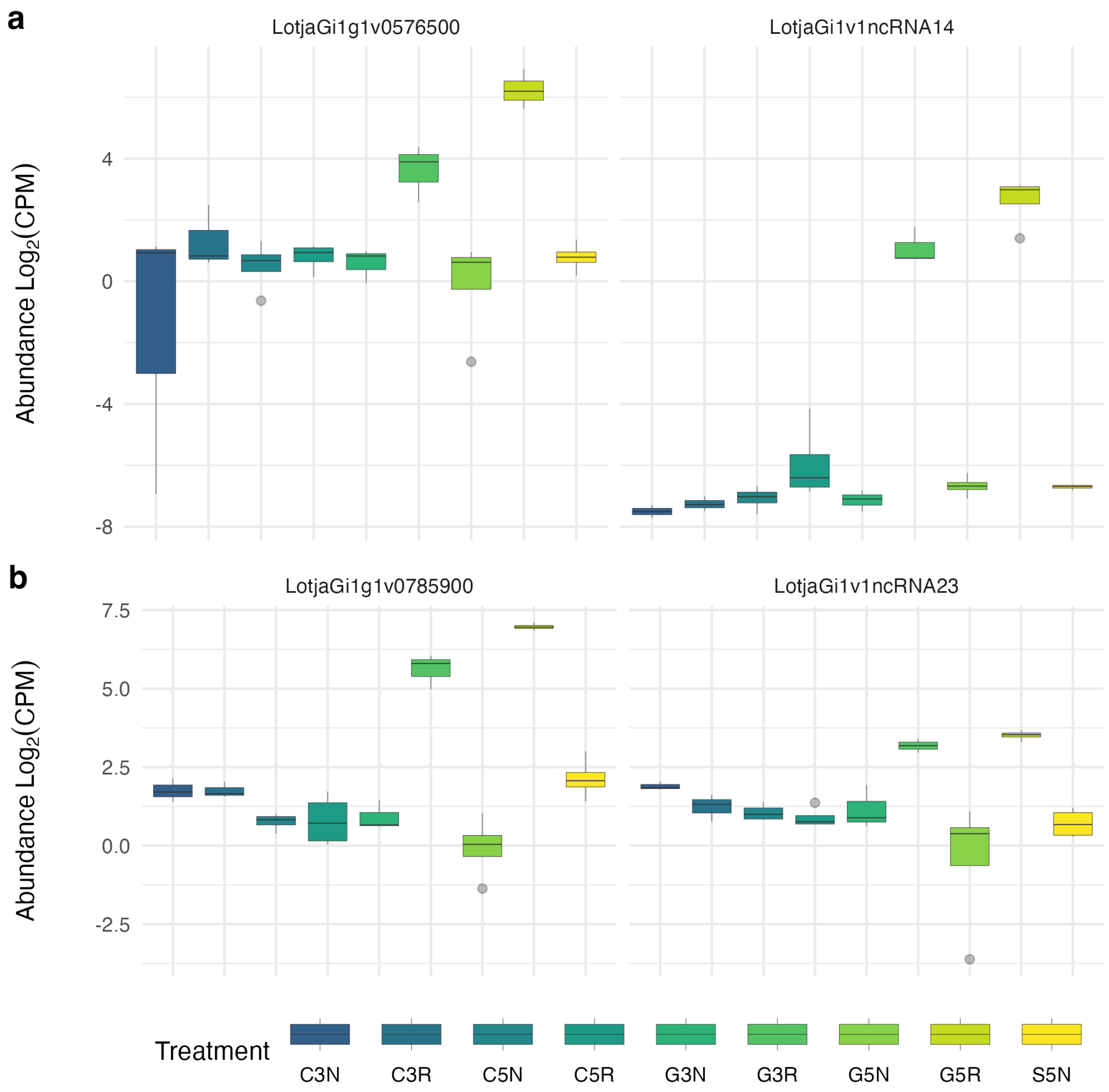


**Fig. S7.** Boxplots of abundance of two symbiosis upregulated ncRNAs and local antisense genes. (A) The proximal gene LotjaGi1g1v0576500, a putative universal stress protein gene (left), and the novel noncoding RNA LotjaGi1v1ncRNA14 (right). (B) The proximal gene LotjaGi1g1v0785900, an F-box protein interaction domain protein (left), and the novel noncoding RNA LotjaGi1v1ncRNA23 (right). N=3 for C3N, C3R, G3N, G3R and N=4 for all other treatments.

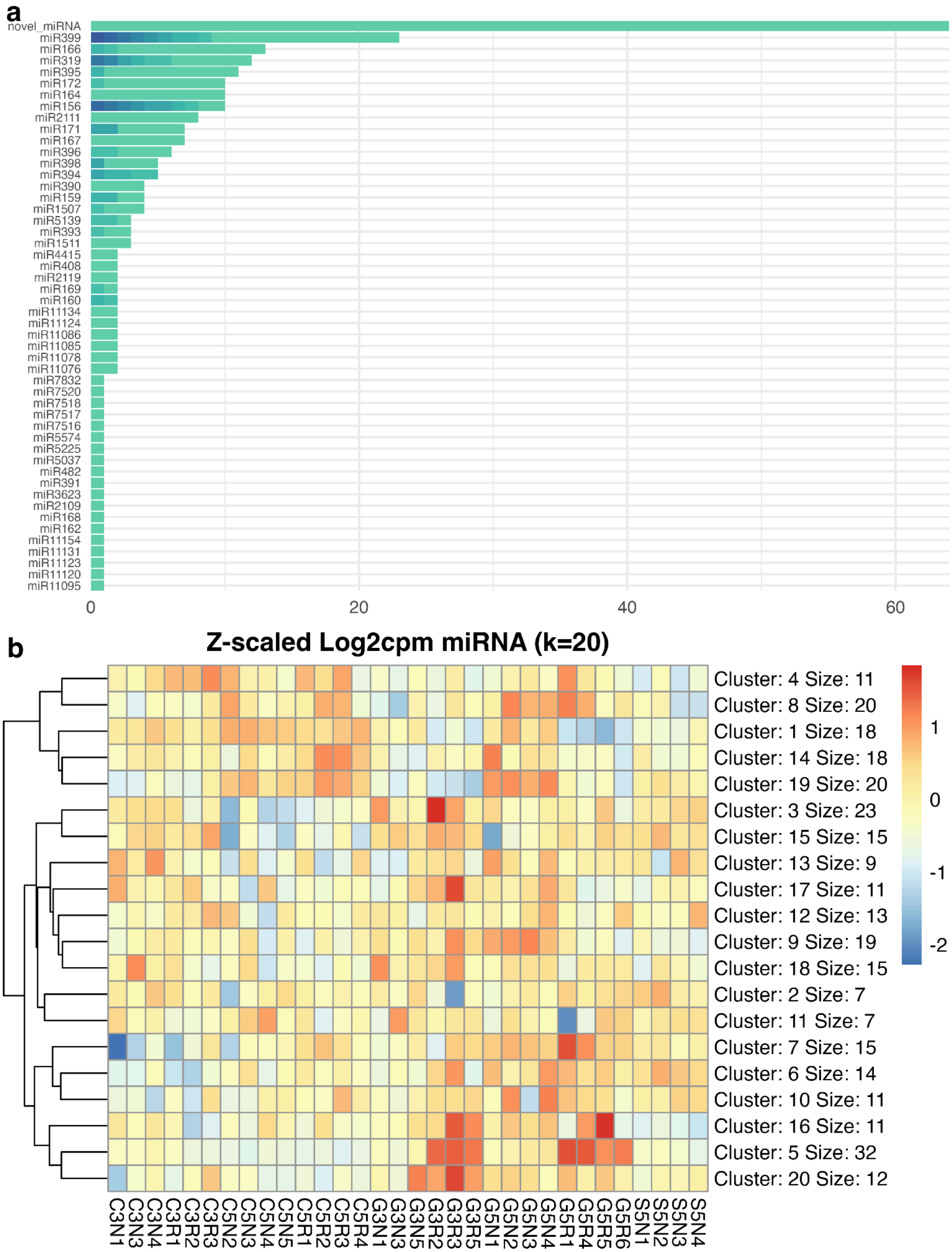
a
novel_miRNA
miR399
miR166
miR319
miR395
miR172
miR164
miR156
miR2111
miR171
miR167
miR396
miR398
miR394
miR390
miR159
miR1507
miR5139
miR393
miR1511
miR4415
miR408
miR2119
miR169
miR160
miR11134
miR11124
miR11086
miR11085
miR11078
miR11076
miR7832
miR7520
miR7518
miR7517
miR7516
miR5574
miR5225
miR5037
miR482
miR391
miR3623
miR2109
miR168
miR162
miR11154
miR11131
miR11123
miR11120
miR11095
0
20
40
60
b
Z-scaled Log2cpm miRNA (k=20)
Cluster: 4 Size: 11
Cluster: 8 Size: 20
Cluster: 1 Size: 18
Cluster: 14 Size: 18
Cluster: 19 Size: 20
Cluster: 3 Size: 23
Cluster: 15 Size: 15
Cluster: 13 Size: 9
Cluster: 17 Size: 11
Cluster: 12 Size: 13
Cluster: 9 Size: 19
Cluster: 18 Size: 15
Cluster: 2 Size: 7
Cluster: 11 Size: 7
Cluster: 7 Size: 15
Cluster: 6 Size: 14
Cluster: 10 Size: 11
Cluster: 16 Size: 11
Cluster: 5 Size: 32
Cluster: 20 Size: 12
1
0
-1
-2
C3N1
C3N3
C3N4
C3R1
C3R2
C3R3
C5N2
C5N3
C5N4
C5N5
C5R1
C5R2
C5R3
C5R4
G3N1
G3N3
G3N5
G3R2
G3R3
G3R5
G5N1
G5N2
G5N3
G5N4
G5R1
G5R4
G5R5
G5R6
S5N1
S5N2
S5N3
S5N4

**Fig. S8.** Full graphs of miRNA family count and miRNA expression K-means clustering. (A) Number of miRNA genes identified across different miRNA families, a color gradient is assigned to each lettered version of a miRNA gene identified (i.e. miR156c, miR156d, etc.) as well as the unlettered version (i.e. miR156). Corresponds to Fig. 4a in the main text. (B) Heatmap of k-means clusters of z-scaled miRNA expression ($Log_2$ Counts Per Million Mapped Reads).

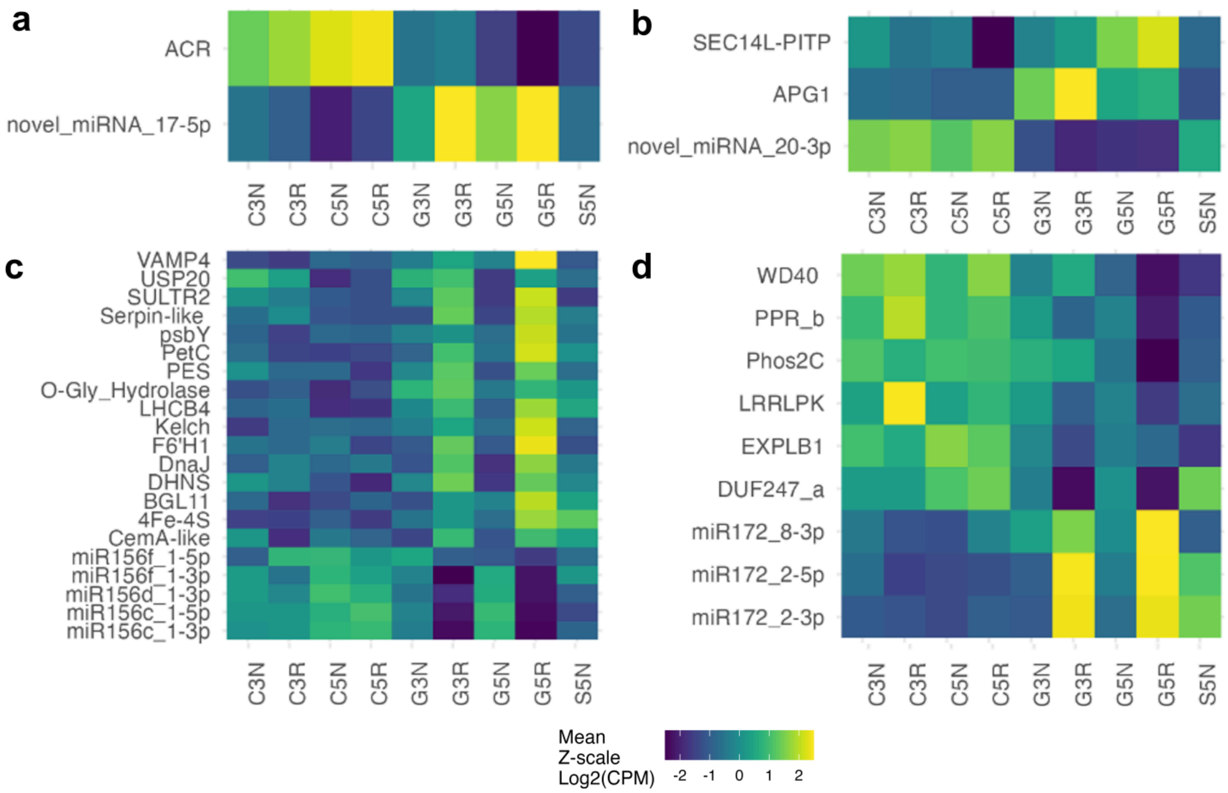


**Fig. S9.** Expression of select miRNAs and correlated targets. Expression is shown as treatment average Z-scaled log2(CPM) of select miRNAs and targets. All miRNAs belong to either cluster 1 (symbiosis downregulated) or 5 (symbiosis upregulated). All displayed miRNAs and predicted target genes have significant negative correlations (Pearsons, FDR <0.05) to at least one member in the same plot. Plots are grouped by miRNA family: (A) novel novel_miRNA_17-5p, (B) novel_miRNA_20-3p, (C) miR156, (D) miR172.

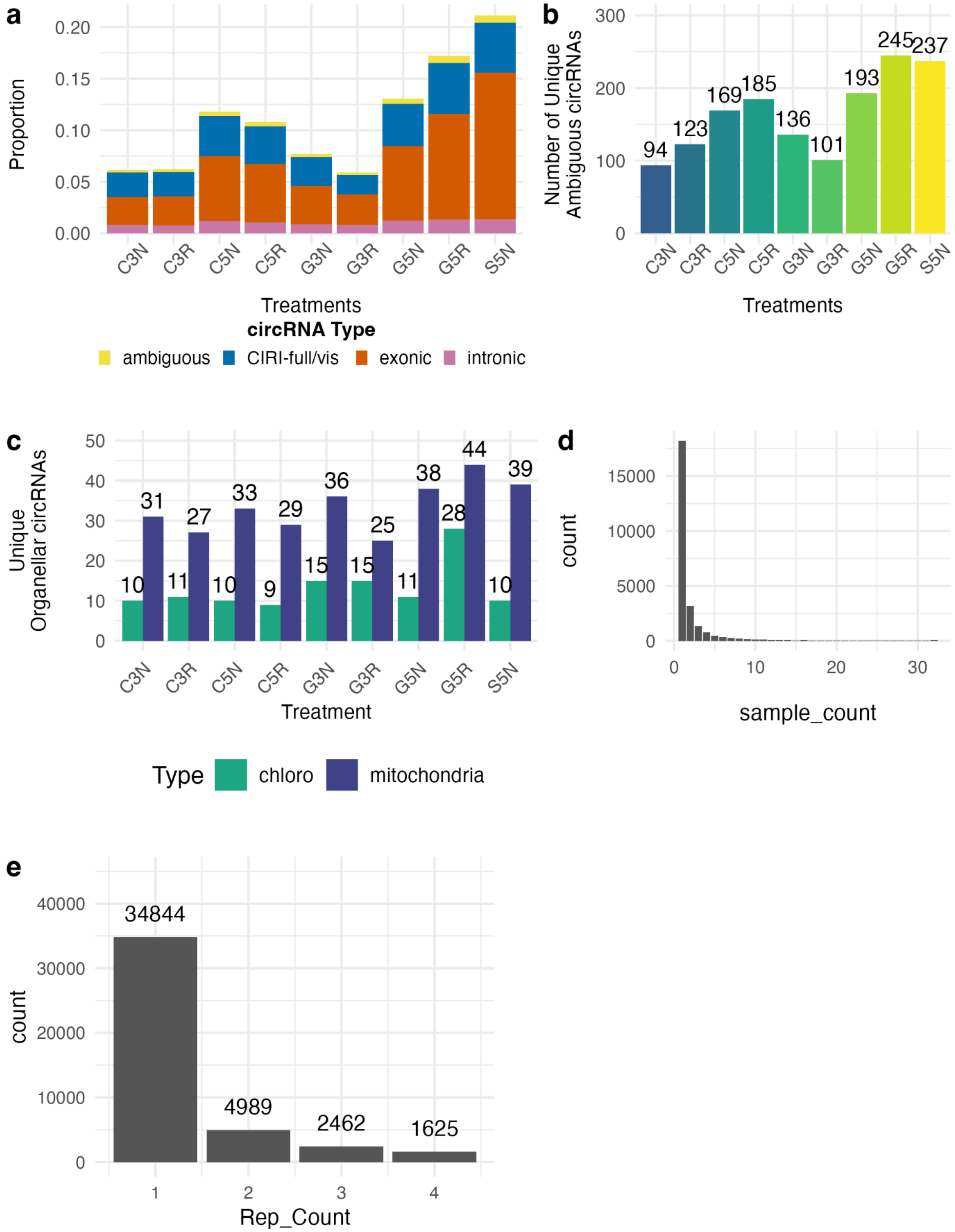
a
Proportion
0.20
0.15
0.10
0.05
0.00
C3N C3R C5N C5R G3N G3R G5N G5R S5N
Treatments
circRNA Type
ambiguous CIRI-full/vis exonic intronic
b
Number of Unique
Ambiguous circRNAs
300
200
100
0
94 123 169 185 136 101 193 245 237
C3N C3R C5N C5R G3N G3R G5N G5R S5N
Treatments
c
Unique
Organellar circRNAs
50
40
30
20
10
0
10 31 11 27 10 33 9 29 15 36 15 25 11 38 28 44 10 39
C3N C3R C5N C5R G3N G3R G5N G5R S5N
Treatment
Type chloro mitochondria
d
count
15000
10000
5000
0
0 10 20 30
sample_count
e
count
40000
30000
20000
10000
0
34844
4989
2462
1625
1 2 3 4
Rep_Count

**Fig. S10. CircRNA genome and replicate distributions.** (A) Bargraph showing the proportion of circRNAs that mapped to exonic, intronic, or ambiguous loci or were only detected by CIRI-full (which does not assign a loci). The ambiguous category includes circRNAs which map to intergenic loci, overlap with genic and intergenic regions, or were assigned differently between CLEAR and CIRI2. (B) The number of individual circRNAs that were assigned as ambiguous across each treatment. (C) The number of individual circRNAs that were assigned to organellar loci across each treatment. (D) Histogram showing the number of samples each circRNA was detected in, the majority of circRNAs were detected in only a single sample. (E) Histogram showing the number of bioreps circRNAs were detected in. CircRNAs are counted on a per-treatment basis here so a circRNA that is found in all 4 replicates of two treatments would be counted twice in column 4.

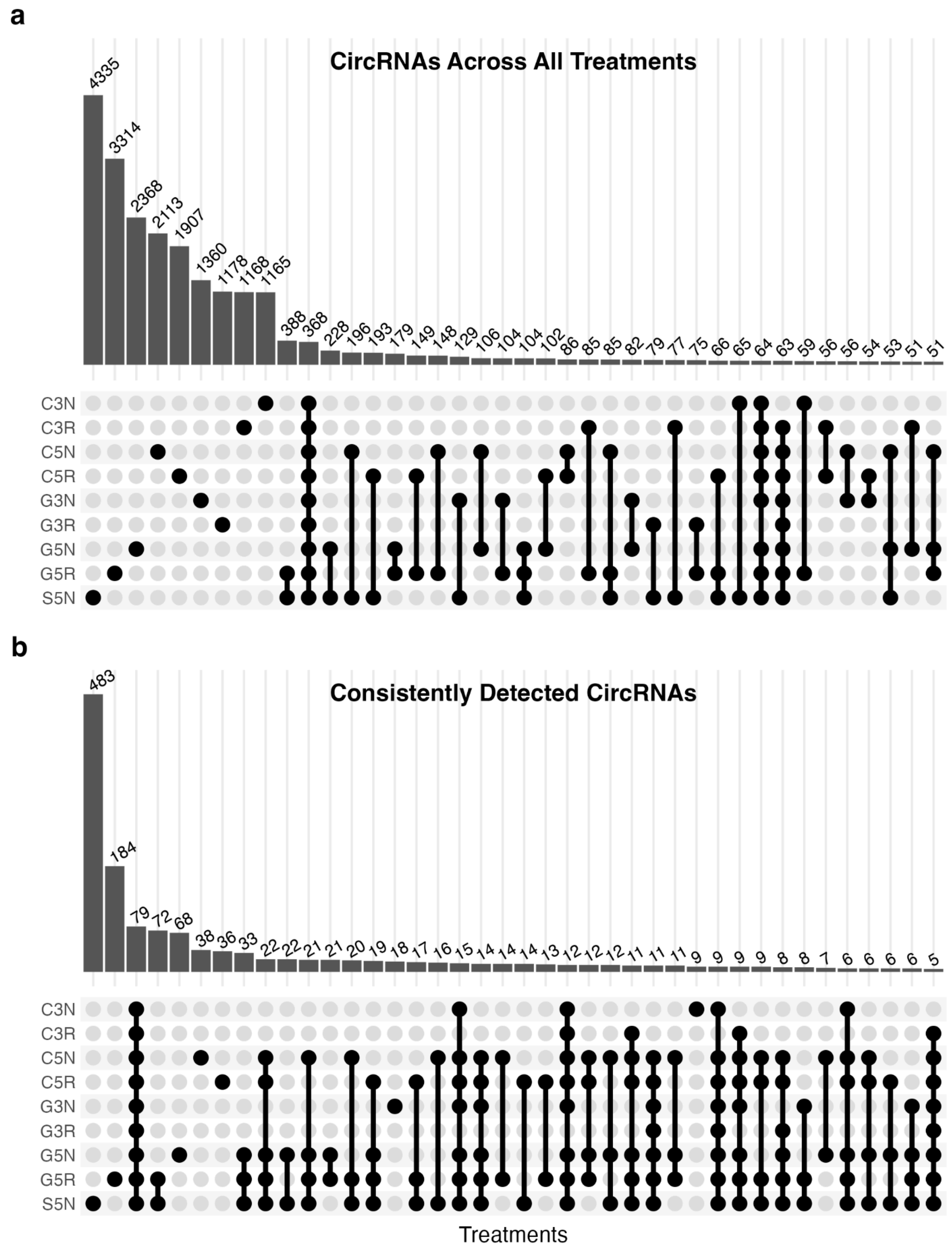


**Fig. S11. CircRNA Presence and Consistency.** (A) Upset plot of all circRNAs observed across treatments. Each individual treatment has more unique circRNAs than any intersection of treatments. S5N+G5R have the most overlap in circRNA observations followed by circRNAs that were observed in all treatments. (B) Full Upset plot of consistently detected circRNAs across treatments from. This upset includes only circRNAs that were in at least 3 replicates of the same treatment. CircRNAs that are in the intersection of multiple treatments are not necessarily in three replicates of multiple treatments, but are in at least three replicates of at least one treatment in the intersection.

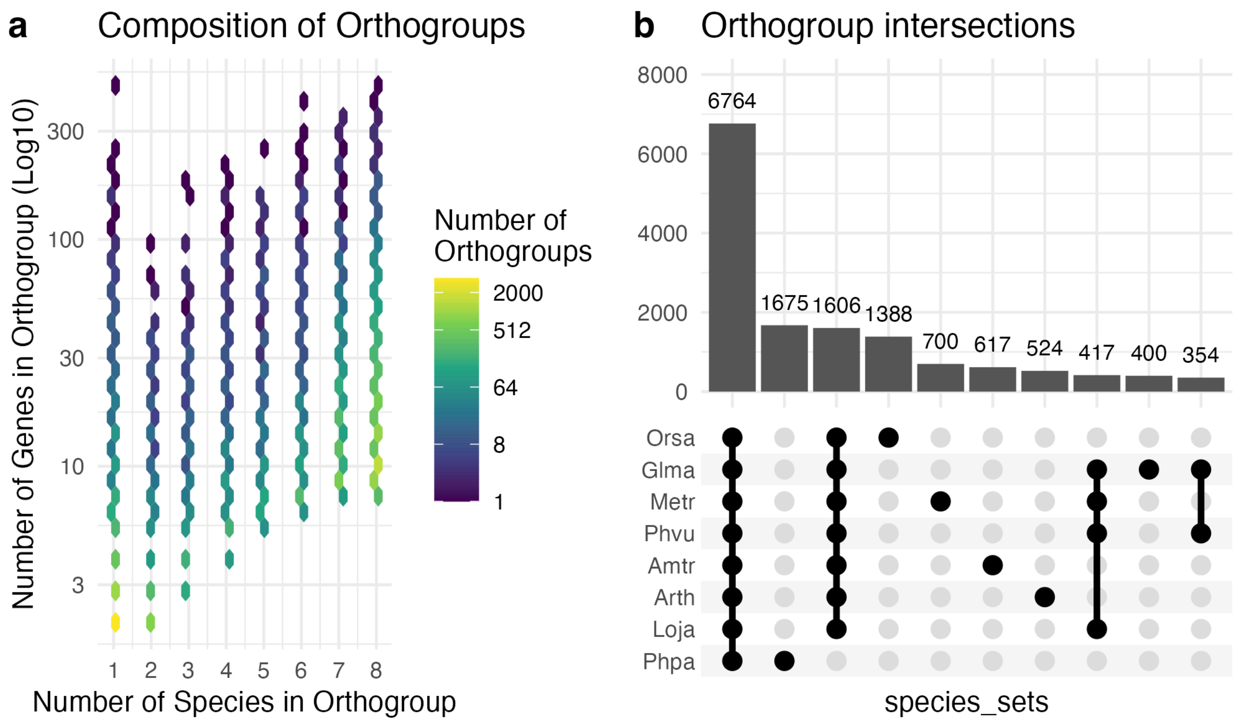


**Fig. S12. Orthogroup composition and intersections.** (A) Binned hexmap showing the number of genes in each orthogroup and the number of species each orthogroup appears in. (B) Upset plot showing the number of orthogroups in the 10 most abundant intersections of species. Amtr = *Amborella Trichopoda* , Arth = *Arabidopsis thaliana*, Glma = *Glycine max,* Loja = *Lotus japonicus,* Metr = *Medicago truncatula,* Orsa = *Oryza sativa*, Phvu = *Phaseolus vulgaris,* Phpa = *Physcomitrium patens*.

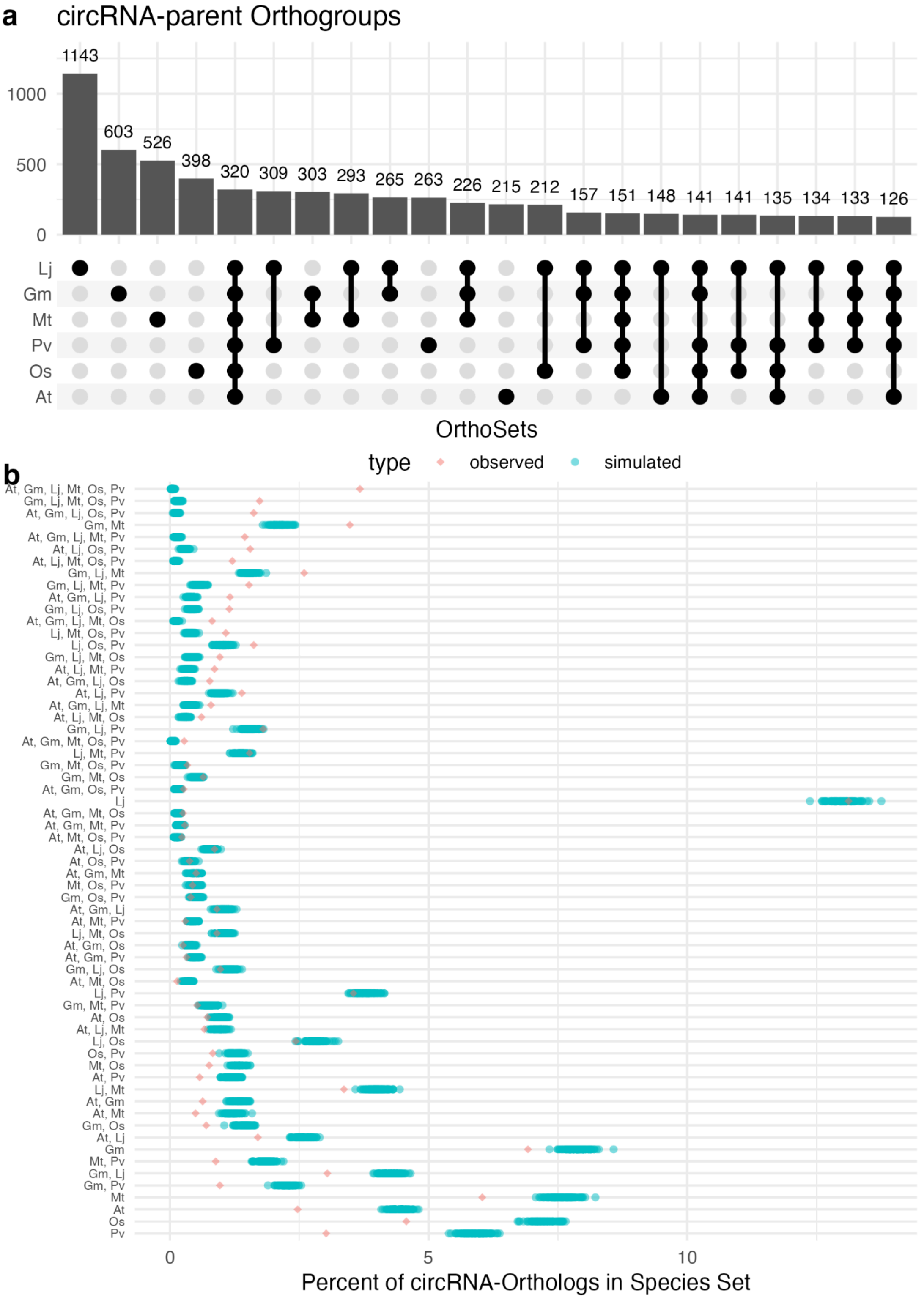


**Fig. S13. Conservation of circRNA parent genes.** (A) Intersections of orthogroup genes which act as circRNA parents in one or more species across the 22 largest intersections (At, *Arabidopsis thaliania;* Gm, *Glycine max;* Lj, *Lotus japonicus;* Mt, *Medicago truncatula*; Os,

*Oryza sativa*; Pv, *Phaseolus vulgaris*). (B) Comparison of observed rates of shared circRNA-parent orthologs across species to simulated rates based on random assignment of circRNAs to genes for each species. Red diamonds to the right of blue points indicate that more common circRNA parents were observed than was expected.

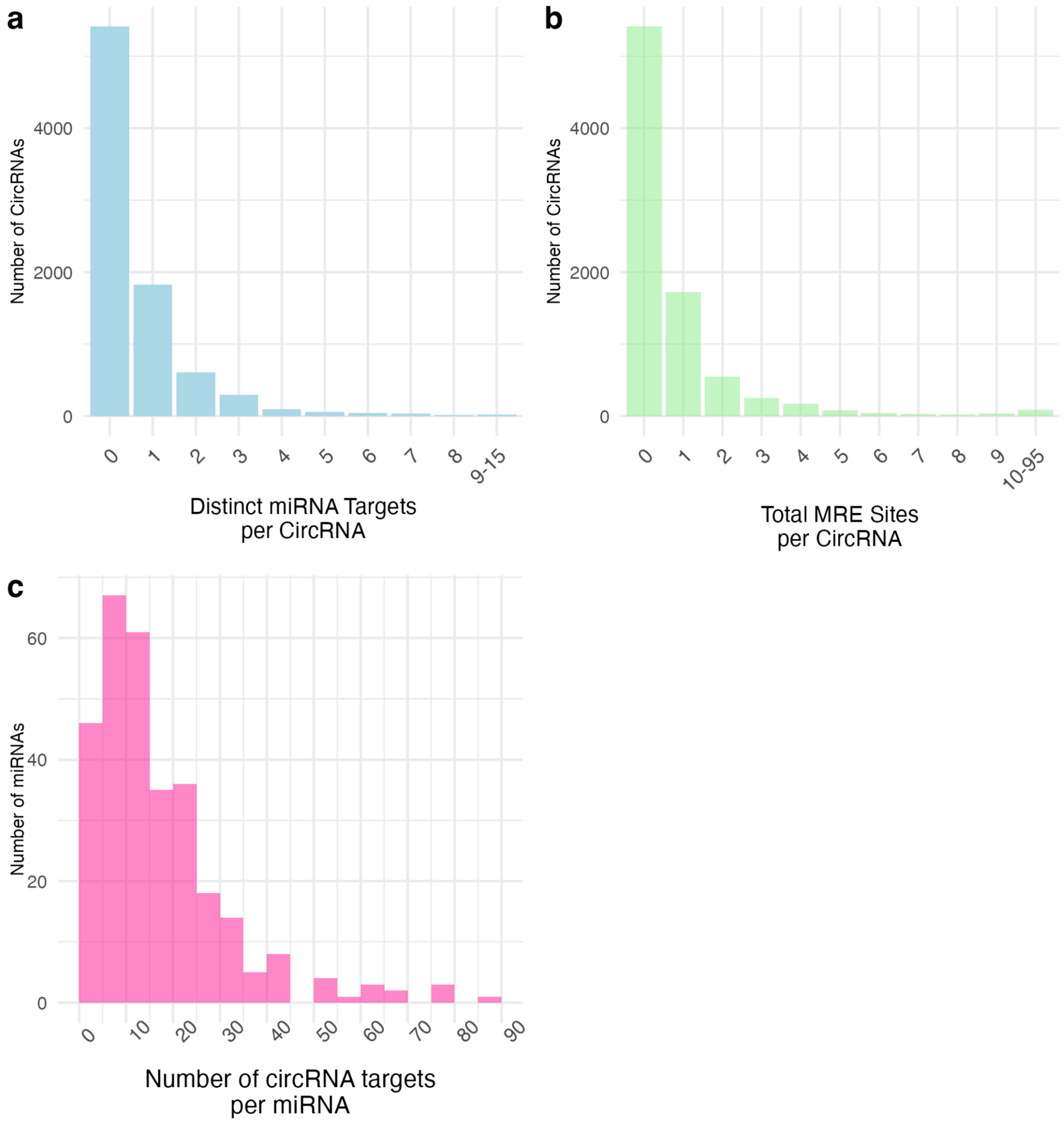


**Fig. S14. MicroRNA targeting of CircRNAs.** (A) Histogram showing the number of distinct microRNAs that circRNAs are predicted to be targeted by. The majority of circRNAs are not predicted to be targets while a small population of circRNAs are targeted by multiple different miRNAs. (B) Histogram showing the distribution of the total number of microRNA response elements on circRNAs. (C) Histogram showing the number of circRNA targets per miRNA.

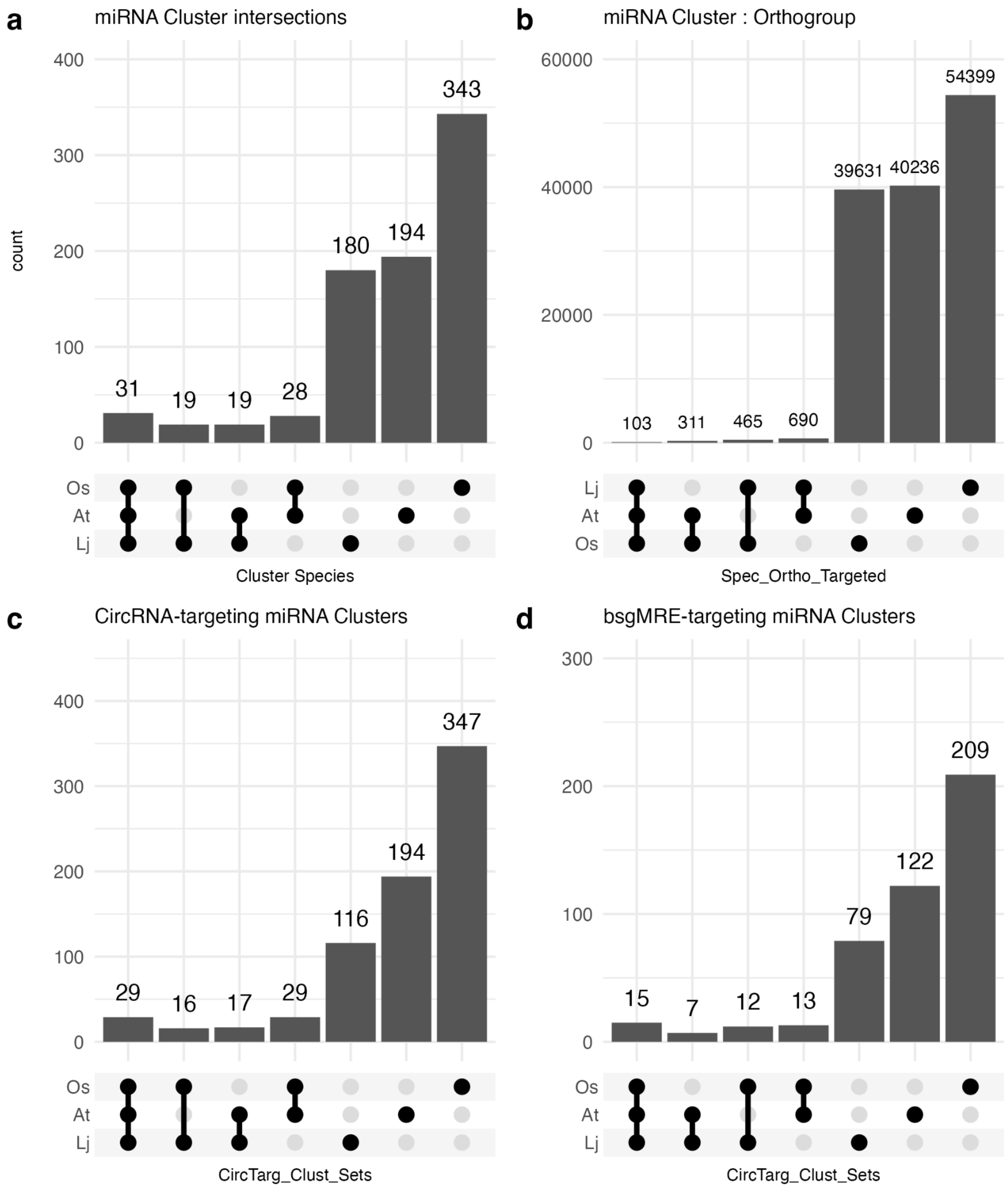

**Fig. S15. MiRNA target conservation.** (A) Upset plot showing the distribution of miRNA clusters shared between *Arabidopsis thaliana* (At), *Lotus japonicus* (Lj), and *Oryza sativa* (Os). (B) Upset plot showing the distribution of orthogroups that were targeted by the same miRNA cluster across the three species. (C) Upset plot showing the distribution of microRNA clusters that targeted at least one circRNA across the three species. (D) Upset plot showing the distribution of microRNA clusters that targeted at least one bsgMRE across the three species.

**Supplemental Table. 1.** CircRNA MRE characteristics of select differentially expressed circRNAs. Cells with red-fill indicate that that miRNA has a bsgMRE on the corresponding circRNA.

| DE circRNA biological group | # of circRNAs with MREs | # of circRNAs with bsgMREs | circRNA ID and Parent Gene | MRE/miRNA | # of linear transcripts targeted by miRNA | cis-regulatory MRE? |
|---|---|---|---|---|---|---|
| Symbiosis upregulated | 0 | 0 | LjG1.1_chr4:2039295-2039535<br>LotjaGi4g1v0016300, (Potassium Transporter) | n/a | n/a | n/a |
| Up in G5R vs 5wk | 2 | 0 | LjG1.1_chr1:7734886-7737488<br>LotjaGi1g1v0054700 (ser/thr kinase) | miR3623_1-3p | 3213 | No |
| | | | LjG1.1_chr4:82574684-82575480<br>LotjaGi4g1v0453800 (BRUTUS) | novel_miRNA_9-3p | 3378 | No |
| *snf-1* regulated | 81 | 5 | LjG1.1_chr1:88664207-88665094<br>LotjaGi1g1v0399500 (Beta-adaptin-like protein) | novel_miRNA_64-5p | 3403 | No |
| | | | | miR396_1-3p | 2552 | Yes |
| | | | | miR162_1-3p | 2016 | No |
| | | | | novel_miRNA_35-5p | 3550 | Yes |
| | | | LjG1.1_chr1:93615759-93616004 | miR319_1-5p | 3271 | No |
| | | | | miR319b_1-5p | 2864 | No |

| | | | | | | |
|---|---|---|---|---|---|---|
| | | | LotjaGi1g1v0440200 (High mobility group protein) | miR319_2-5p | 3134 | No |
| | | | | miR319_5-5p | 2952 | No |
| | | | LjG1.1_chr5:47861736-47863676 LotjaGi5g1v0189300 (Glycolipid transfer protein domain-containing protein | miR5574_1-5p | 3573 | No |
| | | | | miR168_1-5p | 1813 | No |
| | | | | novel_miRNA_4-5p | 3041 | No |
| | | | | miR396_1-5p | 3260 | No |
| | | | | miR396_3-5p | 3345 | No |
| | | | LjG1.1_chr5:9476582-9479080 LotjaGi5g1v0056000 (Protein DETOXIFICATION) | miR5574_1-5p | 3573 | No |
| | | | | miR395_10-5p | 3365 | No |
| | | | | miR395_9-5p | 3159 | No |
| | | | | miR395_5-5p | 3365 | No |
| | | | | miR395_8-5p | 3287 | No |
| | | | LjG1.1_chr6:5839970-5840860 LotjaGi6g1v0031200 (Mediator of RNA polymerase II transcription subunit 1) | novel_miRNA_29-3p | 3396 | No |
| | | | | miR395_2-5p | 2449 | No |

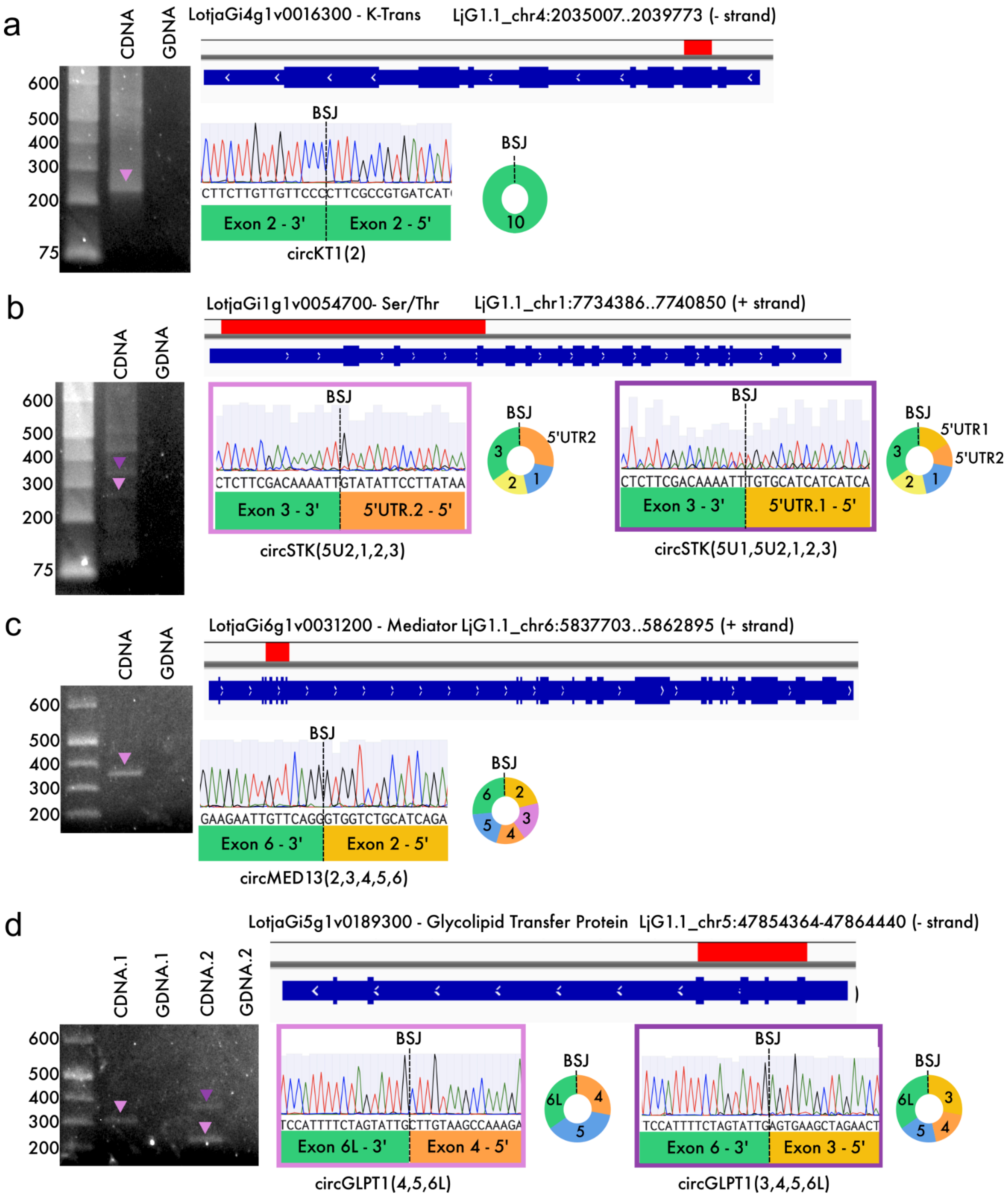
a
CDNA
GDNA
600
500
400
300
200
75
LotjaGi4g1v0016300 - K-Trans
LjG1.1_chr4:2035007..2039773 (- strand)
BSJ
CTTCTTGTTGTTCCCCTTCGCCGTGATCAT
Exon 2 - 3'
Exon 2 - 5'
circKT1(2)
10
b
LotjaGi1g1v0054700- Ser/Thr
LjG1.1_chr1:7734386..7740850 (+ strand)
CTCTTCGACAAAATTGTATATTCCTTATAA
Exon 3 - 3'
5'UTR.2 - 5'
circSTK(5U2,1,2,3)
5'UTR2
CTCTTCGACAAAATTTGTGCATCATCATCA
Exon 3 - 3'
5'UTR.1 - 5'
circSTK(5U1,5U2,1,2,3)
5'UTR1
c
LotjaGi6g1v0031200 - Mediator LjG1.1_chr6:5837703..5862895 (+ strand)
GAAGAATTGTTCAGGGTGGTCTGCATCAGA
Exon 6 - 3'
Exon 2 - 5'
circMED13(2,3,4,5,6)
d
LotjaGi5g1v0189300 - Glycolipid Transfer Protein LjG1.1_chr5:47854364-47864440 (- strand)
CDNA.1
GDNA.1
CDNA.2
GDNA.2
TCCATTTTCTAGTATTGCTTGTAAGCCAAAGA
Exon 6L - 3'
Exon 4 - 5'
circGLPT1(4,5,6L)
TCCATTTTCTAGTATTGAGTGAAGCTAGAACT
Exon 6 - 3'
Exon 3 - 5'
circGLPT1(3,4,5,6L)

**Fig. S16: CircRNA Validation** CircRNAs were validated using divergent primers designed to capture the backsplice junction and to maximize the potential of capturing multiple circs that may have a shared exon. A-D are composed of a locus diagram, a gel image, an excerpt of Sanger sequencing supporting the BSJ, and a diagram of the validated circRNA composition. The locus diagram shows linear transcripts with wide exons and narrow introns in dark blue, and circRNA coordinates identified from NGS data shown as red bars. The gel image shows Divergent primer RT-PCR using cDNA created from total RNA and PCR using gDNA as the template with the same primers. Pink and purple triangles on the gel indicate bands that were gel extracted, and Sanger sequenced, and correspond to the sequencing shown. In the excerpt of Sanger sequencing and the diagram of validated circRNAs, the grey dashed lines show the location of the BSJ. (A) validation of *circKT1(2)* from potassium transporter 1 (LotjaGi4g1v0016300). (B) validation of two circRNAs, circSTK(5U2,1,2,3) and circSTK(5U1,5U2,1,2,3), from Seroine/Threionine (LotjaGi1g1v0054700). (C) validation of two circRNAs, circGLPT1(4,5,6L) and circGLPT1(3,4,5,6L), from Glycolipid (LotjaGi5g1v0189300) and (D) validation of circMED13(2,3,4,5,6) from Mediator (LotjaGi6g1v0031200).

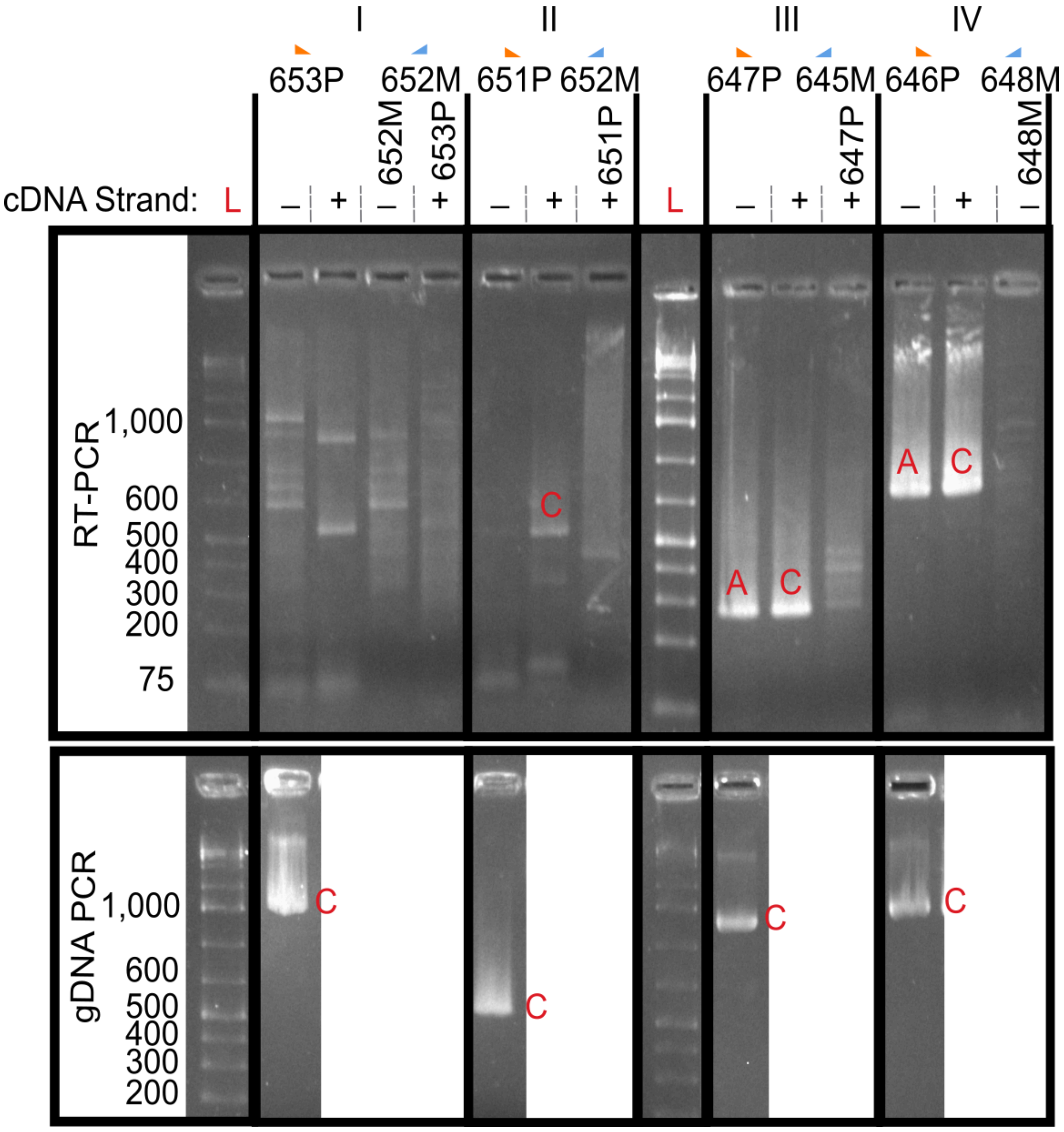


**Fig. S17. BRUTUS Locus Validation:** A) Diagram of BRUTUS (LotjaGi4g1v0453800) locus transcript models. LotjaGi4g1v0453700 is encoded on the + strand and is depicted in orange with two different transcript models; one from LotusBase, and one from Stringtie assembled transcripts generated from our RNAseq data. The Stringtie transcript model has a tail-to-tail overlap with the BRUTUS transcript. The BRUTUS transcript model is shown in blue and is encoded on the - strand. Diagrams of RNA strands from the + and - strands corresponding to the annotated genes are shown along with primer names and locations. Primers ending in P are + strand primers and primers ending in M are - strand primers. B) Gel showing RT-PCR and gDNA PCR amplicons for convergent primer reactions. Gel lanes are grouped by primer sets I through IV which correspond to regions of the locus that were tested for + and - strand transcription. Lanes designated with a red "L" are ladders. Bands designated with a red "C" match the canonical transcript or reference sequence and bands designated with a red "A" are the same sequence but in opposite strand orientation. Region I includes the tail ends of LotjaGi4g1v0453700 and LotjaGi4g1v0453800, neither the + or - strand RT-PCR reactions produced amplicons which aligned to the locus, single primer control reactions where both RT and PCR were performed with only a single primer produced artefacts and primer dimers. The gDNA PCR amplicon in Region I matched the reference sequence. Region II corresponds to the terminal exon and 3' UTR of LotjaGi4g1v0453800, gDNA and + strand cDNA reactions match the expectation of the reference and no evidence of antisense transcription in this region was observed. Region III measures from BRUTUS exon 11 to the terminal exon of BRUTUS. Identical products were observed in both the + and - strand RT-PCR which indicates a natural antisense RNA within the BRUTUS locus. Region III gDNA PCR produced the expected amplicon that matched the reference sequence. Region IV ran from BRUTUS exon 7 to exon 11 and showed amplicons corresponding to both antisense and canonical transcripts with identical sequences. C) Gel showing RT-PCR for divergent primers for + and - strand and with and without RNAse R treatment. Purple and pink triangles indicate bands that were sanger sequenced and correspond to the excerpts of sanger sequence presented in panel D). D) Excerpts of sanger sequence alignments which cover the BSJs and circRNA diagrams for circBRUT(10,11) and circBRUT(10,11,12,13).
