## Supplementary material for "Identification of Potential Regulatory Non-Coding RNAs in *Lotus Japonicus* Symbiosis": SF1 - Supplementary Methods

**Supplemental File 1 - Additional Materials and Methods**

RNA extraction using CTAB protocol (Jordan-Thaden et al., 2015):

Briefly, CTAB extraction buffer (2% (w/v) CTAB, 2% (w/v) polyvinylpyrrolidone K-40, 25mM EDTA, 100mM Tris-HCL, 1g/L spermidine; pH 8.0) with 5% β-mercaptoethanol was pre-warmed at 65 °C, added to the tissue, mixed by vortex, incubated at RT for 5 minutes, and centrifuged at 15,000 × *g* for 10 min. The aqueous phase underwent two sequential 1:1 chloroform:isoamyl alcohol (24:1) extractions before isopropanol precipitation at 4 °C for 1 hr. The RNA pellet was washed twice with 70% ethanol, resuspended in TRIzol (Ambion), and re-extracted with chloroform:isoamyl alcohol, followed by a second isopropanol precipitation. The final pellet was washed with 70% ethanol, dried, and resuspended in 37 °C nuclease-free water. Each sample represented RNA pooled from 15 plants.

Categorization of MSTRGs:

Stringtie assembles RNAseq reads into continuous transcripts, in cases where novel transcripts that do not match a previously annotated isoform are discovered; these transcripts are assigned an "MSTRG" id. In our total RNA sequencing dataset, these "MSTRG" transcripts theoretically include a combination of novel protein-coding genes and isoforms, non-coding RNAs, retrotransposon RNA, fusion genes, and artefacts. We attempted to ensure the accuracy of our gene expression analysis by detecting overlaps between "MSTRG" transcripts and previously annotated protein-coding genes. "MSTRG" transcripts with weak, no, or multiple overlaps with previously annotated genes were left categorized as "MSTRGs" but "MSTRG" transcripts that had >75% overlap with a single annotated gene were re-labeled as coming from that gene for the purposes of gene and isoform expression analysis.

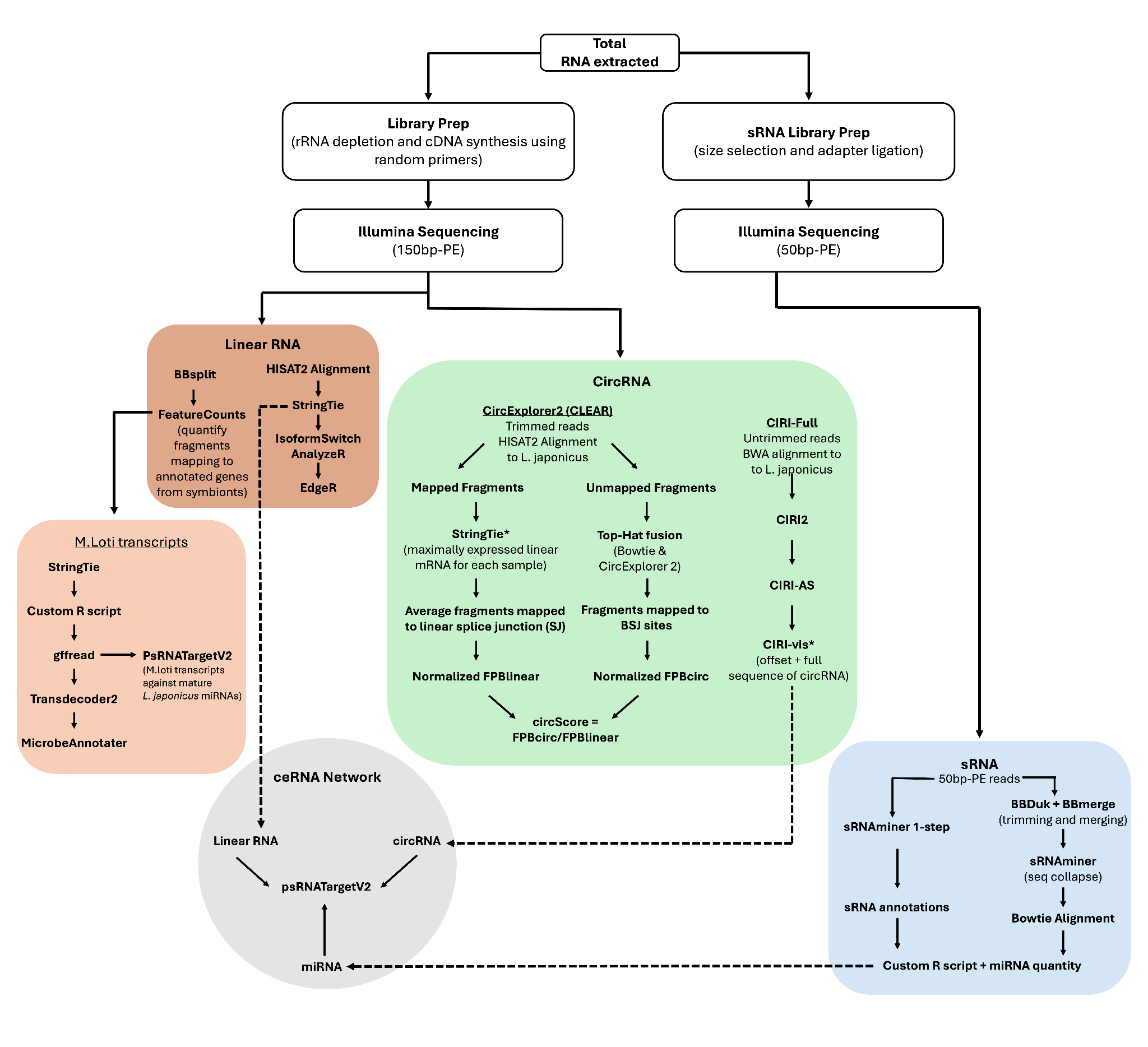

**Supplemental Figure 1: Diagram of bioinformatic workflow.** The same pools of RNA extracted from *L.* *japonicus* root samples underwent small RNA and total RNA-specific library preparation and analysis. Analysis for linear RNAs and circular RNAs is shown in orange and green, respectively, and small RNA analysis is shown in blue. The pipelines results are combined during the analysis of competing endogenous RNA networks.

CircRNA Expression Normalization

Because the two pipelines use different mapping and detection strategies, they produced different read counts for the same circRNAs. To reconcile these differences, we compared read counts from circRNAs detected by both pipelines within the same sample and found them to be tightly correlated (linear regression, R² = 0.96; Supplementary Methods Figure 1). For circRNAs found in multiple samples but not detected by both pipelines, we imputed CIRI read counts into the CLEAR framework using the regression relationship (Supplementary Methods Figure 1), while retaining CLEAR counts when available. The resulting imputed circRNA counts were then used for differential expression (DE) analyses.

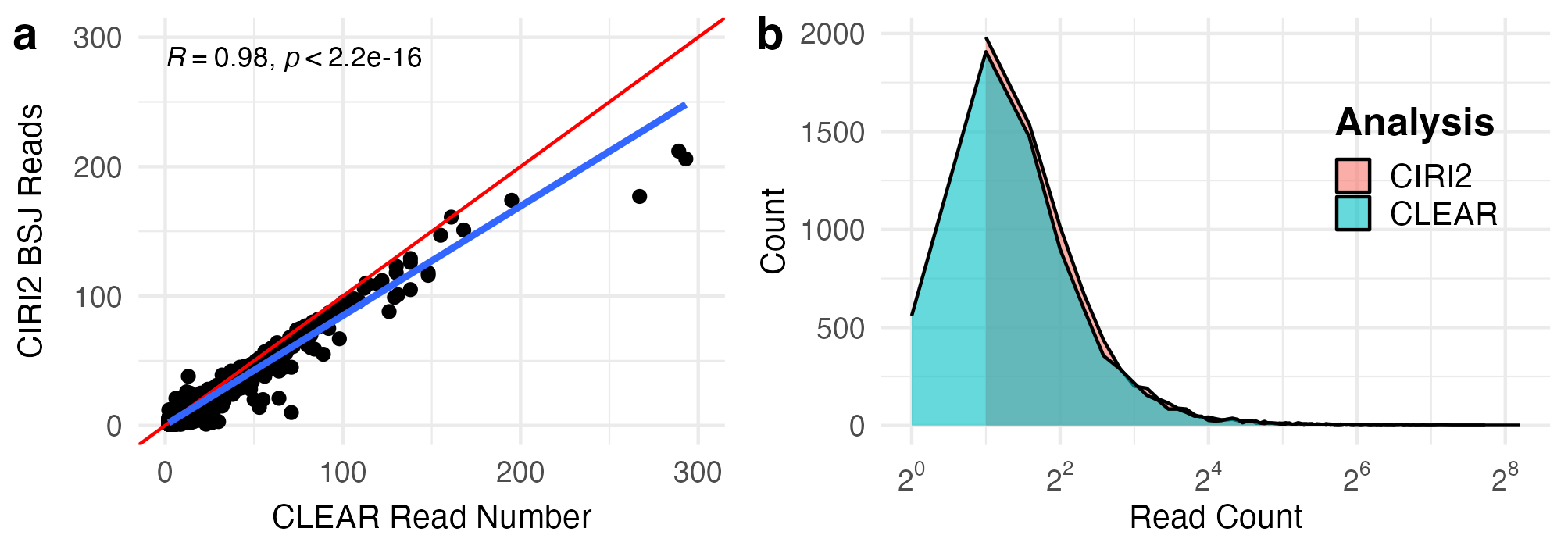

**Supplementary Methods Figure 1. Evidence for Imputed Counts: Correlation of circRNA reads assigned by CIRI2 and CLEAR.** (A) Scatter plot with linear regression of CIRI2 BSJ read counts vs CLEAR junction read counts. Each point shows the read counts from each software for a given observation of a circRNA, and only circRNAs identified by both software in the same replicate are shown. The linear regression line is shown in blue, and a line with a slope of 1 is shown in red. (B) Distribution of read counts from CIRI2 and CLEAR for all circRNAs. The X-axis is log2-transformed for better visibility over the range of the most common read counts. CLEAR includes circRNAs with BSJ counts of 1, whereas CIRI2 requires a putative circRNA to have at least 2 reads spanning the BSJ. Distributions are relatively similar, which also supports imputing reads from CIRI2 into CLEAR for differential expression analysis. Imputed counts can be found in Supplemental File X
